# Selective labelling of arginine residues engaged in binding sulfatedglycosaminoglycans

**DOI:** 10.1101/574947

**Authors:** Thao P. Bui, Yong Li, Quentin M. Nunes, Mark C. Wilkinson, David G. Fernig

**Affiliations:** Department of Biochemistry, Institute of Integrated Biology, University of Liverpool, Crown Street, LiverpoolL69 7ZB, United Kingdom; Liverpool Pancreatitis Research Group, Department of Molecular &Clin. Cancer Medicine, Institute of Translational Medicine, University of Liverpool, 2nd Floor Sherrington Building, Ashton Street, Liverpool L69 3GE, United Kingdom

**Keywords:** arginine, phenylglyoxal, p-hydroxyl phenylglyoxal, fibroblast growth factor, heparansulfate, heparin binding site

## Abstract

The activities of hundreds of proteins in the extracellular space are regulated by binding to the glycosaminoglycan heparan sulfate (HS). These interactions are driven by ionic bonds between sulfate and carboxylate groups on the polysaccharide and the side chains of basic residues in the protein. Here we develop a method to selectively label the guanidino side chains of arginine residues in proteins that engage the anionic groups in the sugar. The protein is bound to heparin (a common experimental proxy for HS) on an affinity column. Arginine side chains that are exposed to solvent, and thus involved in binding, are protected by reaction with the dicarbonyl phenylgyoxal (PGO). Elution of the bound proteins then exposes arginine side chains that had directly engaged with anionic groups on the polysaccharide. These are reacted with hydroxyl-phenylglyoxal (HPG). PGO was found to generate three products: a 1:1 product, the 1:1 water condensed product and a 2:1 PGO:arginine product. These three reaction products and that of HPG had distinct masses. Scripts were written to analyse the mass spectra and so identify HPG labelled arginine residues. Proof of principle was acquired on model peptides. The method was then applied to the identification of heparin binding arginine residues in fibroblast growth factors (FGF) 1 and 2. The data demonstrate that four out of eleven arginine residues on FGF2 and five out of six arginine residues of FGF1 engage heparin. Our approach provides a rapid and reliable means to identify arginines involved in functional complexes such as those of proteins with heparin

## II Introduction

In the extracellular space, interactions between proteins and the glycosaminoglycan heparan sulfate (HS) regulate activities of hundreds of the proteins [1]. These proteins include growth factors, cytokines, chemokines, morphogens, enzymes, enzyme inhibitors, receptors, and extracellular matrix proteins, forming the network of HS-binding proteins, termed the heparan sulfate interactome [2]. Binding to the polysaccharide may, for example, regulate ligand diffusion [3,4], formation of signalling ligand-receptor complexes [5,6] and enzyme activity [7].

The engagement of HS to protein often occurs on the surface or in shallow grooves of proteins with a major contribution from ionic interactions, due to the highly anionic nature of HS. These ionic interactions occur between the negatively charged sulfate and carboxyl groups on the polysaccharide chain with the positively charged residues, lysine, and arginine in proteins [8,9]. Proteins may have more than one HS binding site, and the amino acids contributing to binding though adjacent on protein surface are usually not continuous in the amino acid sequences [10–15].

The structural basis of the interaction of a protein with HS is often probed using the related polysaccharide, heparin, as an experimental proxy. Many approaches such as NMR spectroscopy [16], site-directed mutagenesis [17–19] and X-ray crystallography [6,20], are low throughput. In the absence of a robust bioinformatics method to predict heparin binding sites in proteins, higher throughput experimental methods have been developed. These latter include a method to selectively label lysine residues involved in heparin binding, called “protect and label”, which was coupled with mass spectrometric identification of labelled peptides [11]. This was based on the reaction of N-hydroxysuccinimide (NHS) with the amino side chain of lysine residues and it has been applied successfully to a number of protein-heparin interactions [12,13,21] and also to an electrostatically driven protein-protein interaction [22]. An interesting feature of this method is that it identifies canonical, high-affinity heparin binding sites in proteins, as well as lower affinity secondary binding sites [12,13,21].

NHS cannot react with the guanidino side chains of arginine residues, yet these are at least as important as lysine side chains in mediating the interactions of proteins with anionic partners such as sulphated GAGs [8]. Moreover, the heparin binding sites of some proteins have arginines, but no lysine residues, e.g., fibroblast growth factor FGF22 (Uniprot ID: Q9HCT0-1). Since the heparin binding sites (HBS) on proteins consist of residues not necessarily adjacent in sequence, the identification of arginines, as well as lysines engaged with heparin would provide a comprehensive picture of the ionic interactions.

The modification of arginine residues in proteins is a challenge, because the guanidino functional group of arginine has a pKa between 11.5 ~ 12.5, which makes it the most basic side chain in a protein and a poor nucleophile. There are limited studies on the chemical modification of arginine residues in proteins and the majority of them rely on the reaction of vicinal dicarbonyl compounds with the guanidino group to form cyclic adducts [23]. The ability of phenylglyoxal (PGO) to selectively modify the guanidino group of arginine was first discovered by Takahashi [24], and has been utilized since, especially in studies of enzyme activity [25–27]. In addition, it has been confirmed that PGO does not react with the α-NH_2_group of lysine, indicating that this reagent selectively modifies the guanidino side chain rather than primary amines [28]. The reaction of PGO with the guanidino side chain of arginine is a quantitativereaction but produces several products which may have contributed to the lack of popularity of this approach. However, modern mass spectrometry combined with automated analysis of spectra should allow the deconvolution of multiple reaction products.

Therefore, we have used PGO and hydroxyl-phenylglyoxal (HPG) to develop a method for selective labelling of arginines that are directly involved in binding heparin. PGO was reacted with the arginine residues that are not involved in binding to heparin and so protect them, then HPG was used to selectively label arginine residues in binding sites. The PGO-arginine reaction has three products, and these were readily distinguishable by mass spectrometry. The method was initially tested on the model of peptides and then on two FGFs that have extremely well characterised heparin binding sites (HBSs), FGF1 and FGF2. The data demonstrated the ability of PGO/HPG to quantitatively and selectively modify the guanidino group of arginine residues on proteins. The arginine residues in the primary HBS of FGF1 and FGF2 were all selectively labelled by HPG. The selectively labelling by HPG of arginines in the secondary binding sites of FGF1 and 2 provides a full structural definition of ionic bonding of these sites to the polysaccharide. Interestingly, the data demonstrated the potential for competition by arginine residues on the HBS3 of FGF1 for the binding to HS and the FGF receptor (FGFR) tyrosine kinase [5].

## III Materials and Methods

### Heparin-binding proteins

Recombinant human FGF1 (UniProt accession P05230 – residues 1-155) and FGF2 (UniProt accession P09038 – residues 134-288) were expressed in C41 Escherichia coli cells using the pET-14b system (Novagen, Merck, Nottingham, UK) and purified, as described previously [29].

### SDS-PAGE and Silver stain

Samples were separated on 15% (w/v) polyacrylamide-SDS gels. Silver staining was done according to Heukeshoven and Dernick[30].

### Selective protection and labelling of arginine side chains in HBSs of proteins using PGO and HPG (Fig 1A)

A step by step guide is available at *protocol.io* at the following link: https://www.protocols.io/view/selective-protection-and-labelling-of-arginine-lys-qqmdvu6

**Figure 1:**
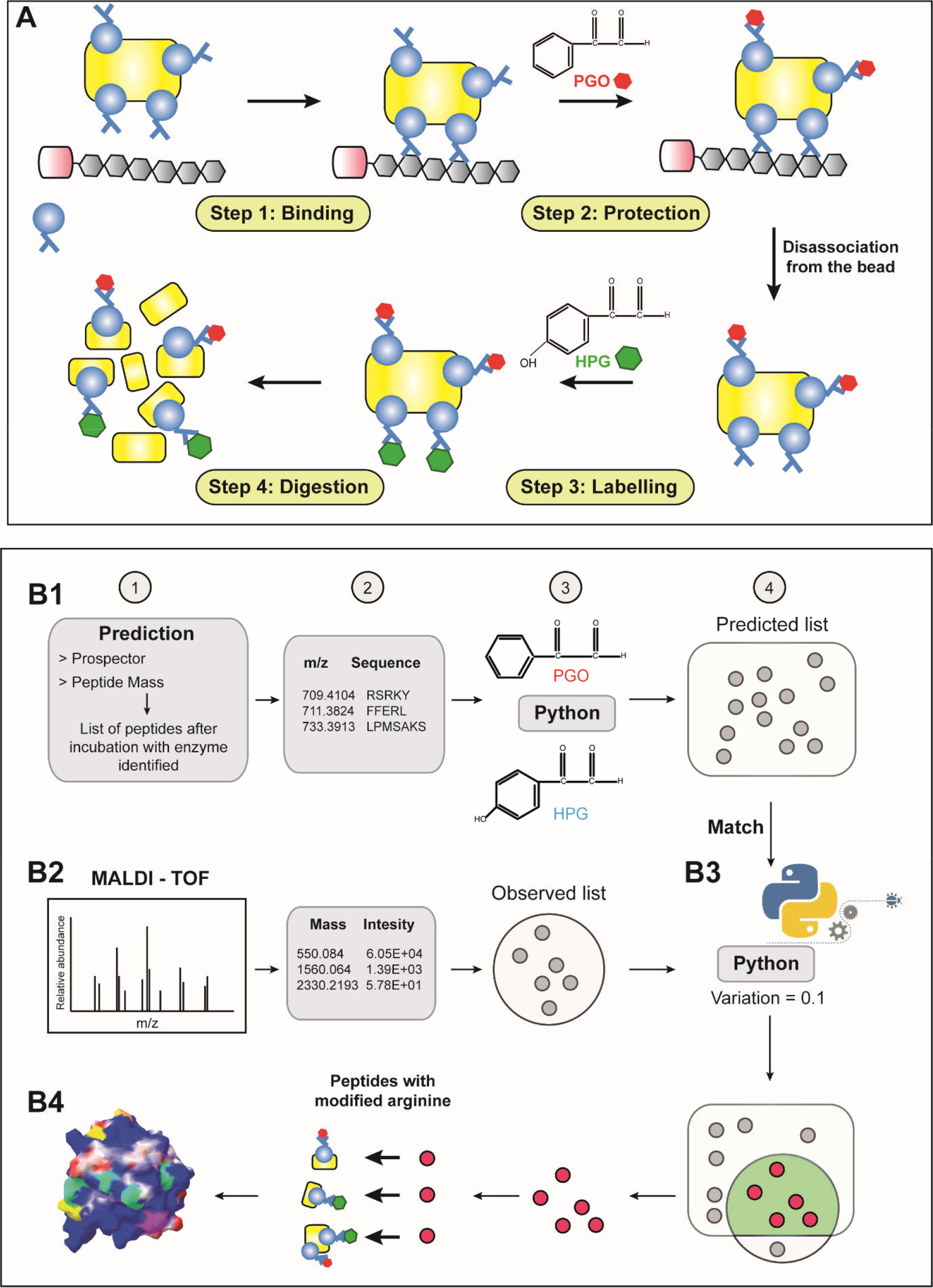
Workflow for selective labelling of arginine residues in proteins. **A. The arginine protect and label experimental workflow**. There are four main steps in the workflow. In the first step, the engagement between protein and heparin affinity beads results in the exposure to solvent of only the non-binding-involved arginine residues. The second step is protection by PGO of these exposed arginine side chains. The protein is then disassociated from heparin with 2 M NaCl, so that the arginine residues in heparin binding sites are available for labelling with HPG in the third step. Finally, proteins are digested by enzymes and peptides are analysed by MALDI-Q-TOF mass spectrometry. **B Analytical workflow for the identification of modified peptides: B1-1, 2.** The corresponding mass and sequence of peptides after enzyme digestion were predicted by Prospector and Peptide Mass. **B1-3.** The peptides were then processed by the *PGO-HPG mass predictor* Python script, which added the possible mass shifts after modification by PGO or HPG on each peptide based on the number of arginine residues in the native sequence. **B1-4.** This step provides the theoretical list of modified peptides. **B2.** The second step starts with the list observed list of peptides from MALDI-TOF MS data, which contains the information of the m/z and intensity of each peak. **B3.** Components in the predicted and observed lists are then matched by the *Matchmaker* Python script with a tolerance of difference of less than 0.1 Da. **B4.** The modified arginine residues were mapped onto the 3D structure of proteins back to identify their locations.

#### Step 1: Binding

AF-heparin beads (Tosoh Biosciences GmbH, Stuttgart, Germany; binding capacity of 4 mg antithrombin III/mL) previously used in the lysine selective labelling protocol [11] were employed. A mini affinity column was made by placing a plastic air filter as a frit at the end of a P10 pipette tip (Star Lab Ltd., Milton Keynes, UK) and then packed with 20 μL AF-heparin beads. The mini-column was equilibrated with 4 × 50 μL 200 mMNaCl, 0.2 M NaHCO_3_, pH 9.5 (Na-1 buffer). The buffer was dispensed slowly into the column using a 2 mL sterile syringe. A minimum of 10 μg FGF protein was loaded onto the column (generally, the loading capacity of FGFs to resin was estimated at 15 mg/mL). The loading was repeated 3 times to ensure the binding between protein and heparin beads. After binding, the column was washed with 200 μL (4 × 50 μL) Na-1 buffer to remove any unbound protein.

#### Step 2: Protection of arginine side chains

PGO (Merck Ltd., UK, 97%) was used in the dark, as it is light sensitive. PGO was freshly prepared in 50 % (v/v) DMSO, 50 % (v/v) HPLC grade water at 1 M, which was then diluted to 0.5 M and then 0.2 M with 0.2 M NaHCO_3_, pH 9.5. The pH was adjusted with 0.1 M NaOH to between 9.1 and 9.5 to ensure optimal reaction. The FGF loaded heparin mini column was rinsed with 30 μL 0.2 M PGO solution to exchange buffers. A further 30 μL PGO solution was added to the column and the bound protein was allowed to react for 60 min at room temperature in the dark. The reaction was quenched with 5 μL 0.1 % (v/v) trifluoroacetic acid (TFA) in water. The mini-column was then washed with 200 μL Na-1 buffer (4 × 50 μL). Bound proteins were eluted with 2 × 20 μL Na-2 buffer (2 M NaCl, 0.2 M NaHCO_3_, pH 9.5) containing 0.1 % (w/v) RapiGest SF Surfactant (Waters, UK). The addition of surfactant was important to ensure protein recovery in this and subsequent steps, due to the increased hydrophobicity of proteins following PGO conjugation to arginine side chains.

#### Step 3: Labelling of Arginine side chain by HPG

The preparation of HPG was performed in the dark room, as it is even more light-sensitive than PGO, following a procedure identical to that used for PGO. The eluted protein was diluted with 400 μL 0.2 M NaHCO_3_, pH 9.5 and concentrated on a 3.5 kDa MWCO centrifugal filter (Merk Millipore, UK) to a final volume of 70~80 μL. The reaction with HPG was performed by incubating 80 μL diluted protein with 20 μL 0.5 M HPG so that the final concentration of HPG in the reaction was 0.1 M. The pH was maintained at over 9.0. The reaction was performed for 60 min at room temperature in the dark and then was quenched with 5 μL 0.1 % (v/v) TFA in water.

#### Step 4: Sample preparation for Mass Spectrometry

Protein was buffer-exchanged by four cycles of dilution on 3.5 kDa MWCO centrifugal filters with 400 μL 10-fold diluted 0.2 M NaHCO_3_, pH 9.5 containing 0.1 % (w/v) RapiGest and 3 cycles of dilution with 400 μL HPLC water containing 0.1% (w/v) RapiGest by centrifugation at 13200 rpm for 10 min. After freezing at −80 °C for 30 min, the sample was lyophilized for an hour.

#### Step 5: Incubation with proteases

##### Chymotrypsin/trypsin

The freeze-dried protein was dissolved in a mixture of 80 μL 25 mM NH_4_HCO_3_ and 10 μL 1 % (w/v) RapiGest (~ 0.1 % w/v in final solution) and heated at 80 ^°^C for 10 min. The mixture was quickly centrifuged at 3200 rpm for 30 seconds before 5 μL 50 mM DTT was added (5 mM final concentration) and incubated for 15 min at 56 °C. After cooling the sample to room temperature, proteins were carbamidomethylated with 5 μL 0.1 M iodoacetamide (freshly made) for 30 min in the dark. Proteins were then digested overnight with chymotrypsin (Promega Ltd., UK) or trypsin (Sigma, UK) at a ratio of 1:100 (w/w).

#### Incubation with Arg-C

The dried sample was dissolved in 8 M urea, 400 mM NH_4_HCO_3_, pH 7.8 (25 μL) and 45 mM DTT (2.5 μL), and incubated for 15 min at 56 °C. After cooling to room temperature, 75 μL incubation buffer (50 mM Tris-Cl, 5 mM CaCl_2_, 2 mM EDTA, pH 7.8) was added to dilute the urea to 2.0 M. Arg-C protease (Promega, Southampton, UK) was freshly prepared in incubation buffer and then added to the protein solution at a ratio of 1:100 (w/w). Activation buffer 10X (50 mM Tris-Cl, 50 mM DTT, 2 mM EDTA, pH 7.8) was added to give a final concentration of 1X. The mixture was mixed gently and centrifuged briefly before allowing digestion to proceed overnight at 37 ^°^C.

#### Step 6: Mass spectrometry for the identification of peptides

Peptides were concentrated by rotary evaporation to a final volume of 10 μL and desalted using C18 Zip-Tips (Millipore). C18 Zip Tips were first pre-wetted with 2 × 10 μL 100 % (v/v) acetonitrile and then pre-equilibrated with 2 × 10 μL 0.1% (w/v) TFA in water. The peptides were loaded on the Zip Tip, the loading was repeated 7 to 8 times to ensure binding. The Zip Tip was washed with 10 μL 0.1% (w/v) TFA. Finally, the peptides were eluted with 2 μL of 5 mg/mL α-cyano-4-hydroxycinnamic acid (CHCA, > 99 % purity, Sigma) in 50:50 acetonitrile/water with 1 % (v/v) TFA, directly on to a 96 spot MALDI (matrix-assisted laser desorption/ionisation) target plate.

Analyses were performed on a Synapt G2-Si (Waters, Manchester, UK) with MALDI source equipped with a frequency tripled Nd:YAG UV laser (λ = 355 nm), operating at 1 kHz. The spectrum acquisition time was 120 seconds, with 1 second scan rates, laser energy of 150 Au. The MS spectra were extracted by MASSLYNX v.4.1 (Waters, Manchester, UK) with the spectrum range from 500 Da to 4000 Da. The spectra were then processed using automatic peak detection, including background subtraction.

#### Identification of modifications of peptides

Protein Prospector (v 5.19.1 developed at the USCF Mass spectrometry Facility) and Peptide Mass (ExPASy) were used to predict the possible peptides after incubation of protein with an enzyme with the following parameters: enzyme, chymotrypsin or trypsin; maximum missed cleavages, 5; mass range, 500 to 4000 Da; monoisotopic; instrument, MALDI-Q-TOF (Fig 1-B1). The list of peptides after enzyme cleavage was filtered to remove the peptides without arginine residues. Because products from the reaction between PGO/HPG and arginine residue bring different additional masses to the peptides, the prediction of the peptide masses after the modification was achieved using a script, written in Python (version 3.5.3 released on January 17th, 2017, available at http://www.python.org), named “*PGO-HPG mass predictor*” (available at Github in following link: https://github.com/bpthao/PGO-HPG-mass-predictor_2). Based on the number of arginine residues in each peptide, “*PGO-HPG mass predictor*” script considers all possible reaction products of PGO and HPG with arginine, and generates a list of predicted masses of the modified peptides (Fig 1-B2).

*PGO-HPG mass predictor* used two inputs. First, the list of predicted peptides from the native proteins cleaved by enzyme from either Protein Prospector or Peptide Mass (ExPASy). The first file is composed of two columns, the native sequence of the peptide and the corresponding mass. Second, the file of modifications also comprises of two columns, the mass shift of each product and its descriptions. Using a loop*, PGO-HPG mass predictor* automatically adds up all potential mass shifts to each peptide with has arginine residues. All combinations of mass changes were covered. The output file has four columns: 1-the native sequence of the peptides; 2- the original mass; 3-the final mass after modifications; 4-the description of the reaction product(s), to facilitate.

The observed list is the original mass of FGF peptides after modifications (Fig 1-B3), which provided the mass and intensity of each peak. The match between predicted and the observed list was carried out with a second Python script, “Matchmaker”, (available at Github in following link: https://github.com/bpthao/Matchmaker_2) with a mass difference tolerance set to 0.1 Da (Fig 1-B3), as recommend by Mascot.

The output is a list of matching peptides with the native sequence, the predicted mass with modifications, the specific modifications associated with arginine residues and the actual observed mass.

## IV Results and discussion

### PGO reactions with arginine in model peptides

The reaction of PGO with arginine, though highly selective, has been demonstrated to result in several products, influenced by the ratio of reactants and the adjacent amino acids [25,31–34]. Thus, according to Takahashi [24], the ratio of PGO to arginine in the reaction determines the product (Fig 2A). The reaction starts with the formation of an adduct of PGO with the guanidino group (Fig 2A, product {1}), which can then reversibly release a hydroxyl group from glyoxal, resulting in the alternative product (Fig 2A, product {2}). Following this is the reactionof a second PGO molecule (if sufficient reactant) onto either a nitrogen of the guanidino group of arginine (Fig 2A, products {3}) or the carbonyl group of the first PGO, which may reversibly release a hydroxyl group from glyoxal to form the further products (Fig 2A, products {4}). All of the reactions are highly dynamic, in the presence of even or excess of PGO, so a mixture of all products is formed, though in solution, product {2} predominates over product {1}[24].

**Figure 2:**
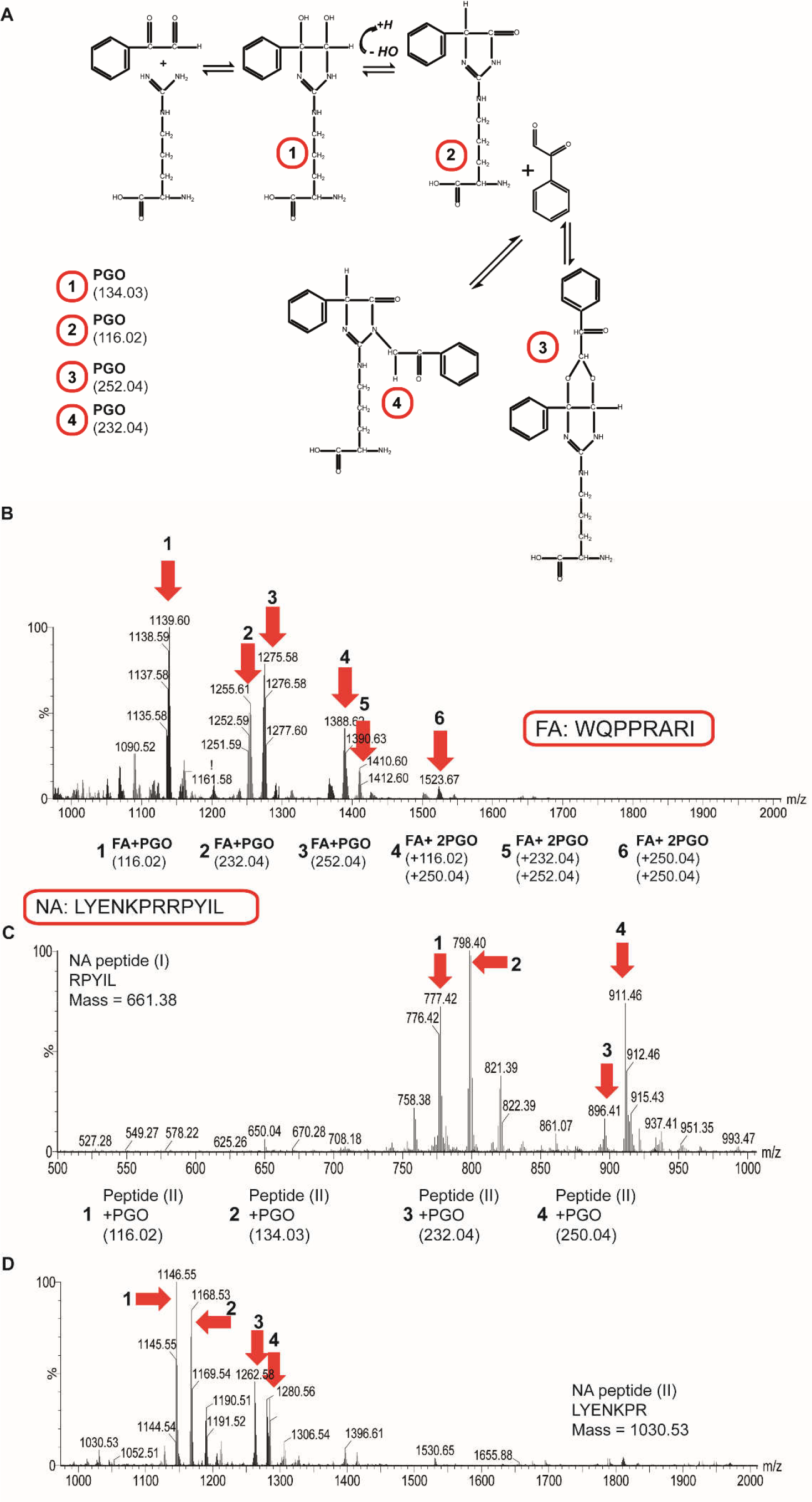
Optimization of the reaction between PGO and arginine residues on peptides. **A. The description of the stoichiometry of PGO with arginine**. Initially, PGO reacts with the guanidine group of arginine to form a first product (Product 1), which may then condens with the loss of a water molecule through a reversible process to yield product 2. The 2:1 water condensed product of PGO and arginine may appear in the presence of excess PGO (product 3 or product 4), which have the same mass. **B. Mass spectra of peptide FA reacted with PGO.** The reaction between FA and 200 mM PGO was conducted in the dark, for 30 min at 25^°^C. There are six products observed. The products between FA and one PGO were demonstrated as three major peaks of 1139.60 Da, 1255.61 Da and 1275.58 Da. The reaction of FA with two PGO results in three peaks, numbered 4, 5 and 6 with the mass of 1388.62; 1410.60 and 1523.67. **C. Reactions of arginine residues located at N- or C-terminus of a peptide.** Peptide NA was cleaved by trypsin to generate two peptides: i) NA-I: RPYIL, mass 661.38 Da with arginine at the N terminus and ii) NA-II: LYENKPR, mass 1030.53 Da with arginine at C terminus. Both peptides were reacted with 200mM PGO in the dark for 30 min. After the reaction, each peptide showed 4 products, indicated by the arrows and numbered 1, 2, 3 and 4.

In contrast, the reaction of HPG with arginine only has one product [35]. Since two reagents are required for selective labelling, one to protect arginines not involved in binding and a second to label arginines engaged in non covalent bonds with the polysaccharide, it was important to not just optimise these reactions, but also to establish which products are formed following reaction with PGO. To determine these parameters, model peptides were used in the first instance. These peptides were specifically chosen to represent arginine residues in particular contexts. Peptide FA (WQPPRARI) has two arginine residues, separated by a single alanine, which will provide insight into any differences in a reaction that may occur due to proximity of arginines on the surface of a protein (Table 1). Peptide NA provides the means to generate an N-terminal arginine (p-Glu-LYENKPRRPYIL) by pre-digestion with trypsin and so identify any influence of the adjacent free amine of the N-terminus and the impact of lysine on PGO reaction with arginine (Table 1B).

**Table 1:** Reactions of arginine residues in peptide FA with PGO. **A. Screen for the conditions for PGO reaction on model peptide FA.** Peptide FA at three different concentrations (100 mM, 200 mM, 1M) reacted with PGO at two different times (5 min and 30 min). (X) corresponds to “does not react” and (V) indicates that the reaction between peptide and PGO was detected. **B. Screen for the products for PGO reaction on model peptide NA.** Peptide NA was pretreated by trypsin to generate two peptides: NA-I and NA-II. The resulting peptides reacted with PGO at concentration of 200 mM for 30 min.

|  | Concentration | 100 mM | 200 mM | 1M |
| --- | --- | --- | --- | --- |
|  | Time |  |  |  |
| **FA**<br>*WQPPRARI*<br>MW:<br>*1023.58 Da* | 5 min | X | V | X |
|  | 30 min | X | V | X |
|  | **PGO 200 mM for 30 min** |  |  |  |
|  | **Products** | **Mass (Da)** |  | **Shift** |
|  | 1 | 1139.60 |  | 116.02 |
|  | 2 | 1255.61 |  | 232.03 |
|  | 3 | 1275.58 |  | 252.0 |
|  | 4 | 1388.62 |  | 116.02+250.04 |
|  | 5 | 1410.60 |  | 232.03+250.04 |
|  | 6 | 1523.67 |  | 250.04+250.04 |

| **Peptide NA**<br>(p-Glu-LYENKPRRPYIL) (PGO200 mM for 30 min) |  |  |  |  |  |
| --- | --- | --- | --- | --- | --- |
| peptide **NA-I**: RPYIL<br>MW: 661.38 Da |  |  | peptide **NA-II**: YENKPR<br>MW: 1030.53 Da |  |  |
| Products | Mass (Da) | Shift | Products | Mass (Da) | Shift |
| **1** | 777.42 | 116.02 | **1** | 1146.55 | 116.02 |
| **2** | 795.40 | 134.03 | **2** | 1164.52 | 134.03 |
| **3** | 896.41 | 232.03 | **3** | 1262.58 | 232.03 |
| **4** | 911.46 | 250.04 | **4** | 1280.56 | 250.04 |

#### The reaction of PGO with arginine residues in peptide FA

First, three concentrations of PGO were tested with peptide FA, which contains two arginines, to identify the conditions required for the reaction to go to completion (Table 1). Two reaction times (5 min and 30 min) were used, each with three different concentrations of PGO. Peptide FA was unmodified after reaction with 100 mM PGO for 5 min or 30 min(Table 1). However, with 200 mM PGO the peak of the original FA peptide was not detected demonstrating that arginines in the peptide were fully reacted with PGO (Fig 2B). No peaks were identified when the peptide was reacted with 1 M PGO, which could be due to the aggregation of the peptide induced by the high concentration of PGO (Table 1). Thus, 200 mM was considered as the optimal concentration of PGO for the modification of arginine in this model peptide.

Interestingly, six reaction products of peptide FA with PGO were detected (Table 1A). The mass of unreacted FA is 1023.58 Da, and after reaction three major peaks of 1139.60 Da, 1255.61 Da and 1275.58 Da were detected, equivalent to a mass shift of 116.02 Da of Product {2} (Fig. 2A) (Peak 1, Fig. 2B, Table 1A); a mass shift of 232.04 Da (Product {4}, Fig. 2A; Peak 2, Fig 2B, Table 1A); and 252.0 Da (Product {3}, Fig. 2A; Peak 3, Fig 2B, Table 1A), respectively.

Peptide FA has two arginine residues and their reaction with PGO also produced three combinations of products. The combinations of product {2} and product {3} resulted in the mass of peak 4of 1388.62 Da (Fig 2B, Table 1A); of product {4} and product {3} resulted inpeak 5 of 1410.60 (Fig 2B, Table 1A); finally, when both arginines formed product {3} with PGO, a mass shift coresponding to peak 6 of 1523.67 was observed (Fig 2B) (Table 1A).

#### The reaction of PGO with argininesin peptide NA

It is clear from the above that the position of arginine resides can affect the reaction product. Since the side chain of an arginine at the terminus of a peptide has more steric freedom, it is more likely that the dicarbonyl group of PGO could form a reversible bond at a 2:1 ratio withthe guanidino group of arginine, as described in Takahashi’s study [24]. To establish if this was the case, two peptides: peptide NA-I (RPYIL, molecular mass 661.38) and peptide NA-II (LYENKPR, molecular mass 1030.53) were produced by treating peptide NA with trypsin (Table 1B). This yielded one peptide with arginine at the N terminus (peptide NA-I) and another one with arginine at the C terminus (peptide NA-II). Both peptides were reacted with 200 mM PGO in the dark for 30 min (Figs 2 C,D).

For peptide NA-I, the product of this peptide with one PGO had amass shift of 116.02 Da (Product {2}, Fig. 1A; Peak 1, Table 1B) resulting in the product of 777.42 Da (Fig 2C), while, the peak at 795.40 Da (Fig 2C) indicated the presence of product {1} from the reaction between arginine and PGO. The addition of a second PGO resulted in the product of 896.41 Da, giving a mass shift of 232.03 Da (Product {4}, Fig 2A, Table 1B) and the product of 911.1 Da due to a mass shift of 250.04 Da (Product {3}, Fig 2A, Table 1B).

In a similar manner, the products of peptide NA-II with PGO showed the mass shifts of all four products between arginine and PGO (Fig 2D, Table 1B). The peak of 1146.55 Da (Table 1B) corresponds to the mass shift of 116.02 Da (Product {2}, Fig 2A); the mass shift due to product {1} (Fig 2A) is detected as the peak at 1164.52 Da (Fig 2D, Table 1B); the product of two PGO reacting with one arginine are observed as the peak at 1262.58 Da (Fig 2D, Table 1B) of product {4} and the peak at 1280.56 (Fig 2D, Table 1B) was due to product {3}. In addition, for the mass shifts observed, only the arginine of peptide II was modified by PGO and there was no reactivity of its lysine or N-terminus towards PGO, in agreement with previous work [28].

The use of peptides with arginine residues in defined contexts demonstrated that 200 mM PGO was likely to fully protect all of the arginine residues in a protein regardless of the adjacent sequence. Moreover, the position of arginine in the peptides affected the formation of the Schiff’s base and types of products produced. However, these are resolvable by mass spectrometry. The next step was to determine whether HPG could similarly fully react with arginine residues.

#### HPG reactions with arginine in model peptides

For the labelling step, HPG, which has the same dicarbonyl moiety as PGO, was chosen. In general, HPG modifies arginine in a similar manner to PGO, but the rate of the reaction is faster than that of PGO and it increases with pH, where pH 9-10 is optimal [35]. HPG forms a single product with the guanidino group of arginine, which is stable at this pH (Fig 3A). The two peptides FA and NA were used again to validate the reactivity of HPG toward to guanidino group of arginine. HPG was dissolved in DMSO at 1 M and diluted stepwise to 500 mM and then 100 mM with 0.2M NaHCO_3_ pH 9.5 and then reacted with the peptides FA and NA for 10 and 30 min. Peptide FA was not entirely modified after reaction with 100 mM HPG for 10 min (Fig 3B), as evidenced by the presence of two major peaks of unmodified and HPG-modified peptide FA, indicating that this time was too short for the reaction to go to completion. However, 100 mM HPG for 30 min modified all of the arginines in the FA peptide (Fig 3B), since the peak of the unmodified peptide FA was no longer detected. In a similar manner, HPG at 100 mM after 10 min reaction did not modify fully the arginines of peptide NA, evidenced by the observation of two major peaks of unmodified and HPG-modified peptide NA, whereas after 30 min, they were fully modified, as only one major peak was seen (Fig 3C). These reactions with the model peptides demonstrated that HPG modifies arginine to form a single product. The difference in the reaction between HPG and arginine compared to that of PGO may be due to the presence of the hydroxyl group in HPG.

**Figure 3:**
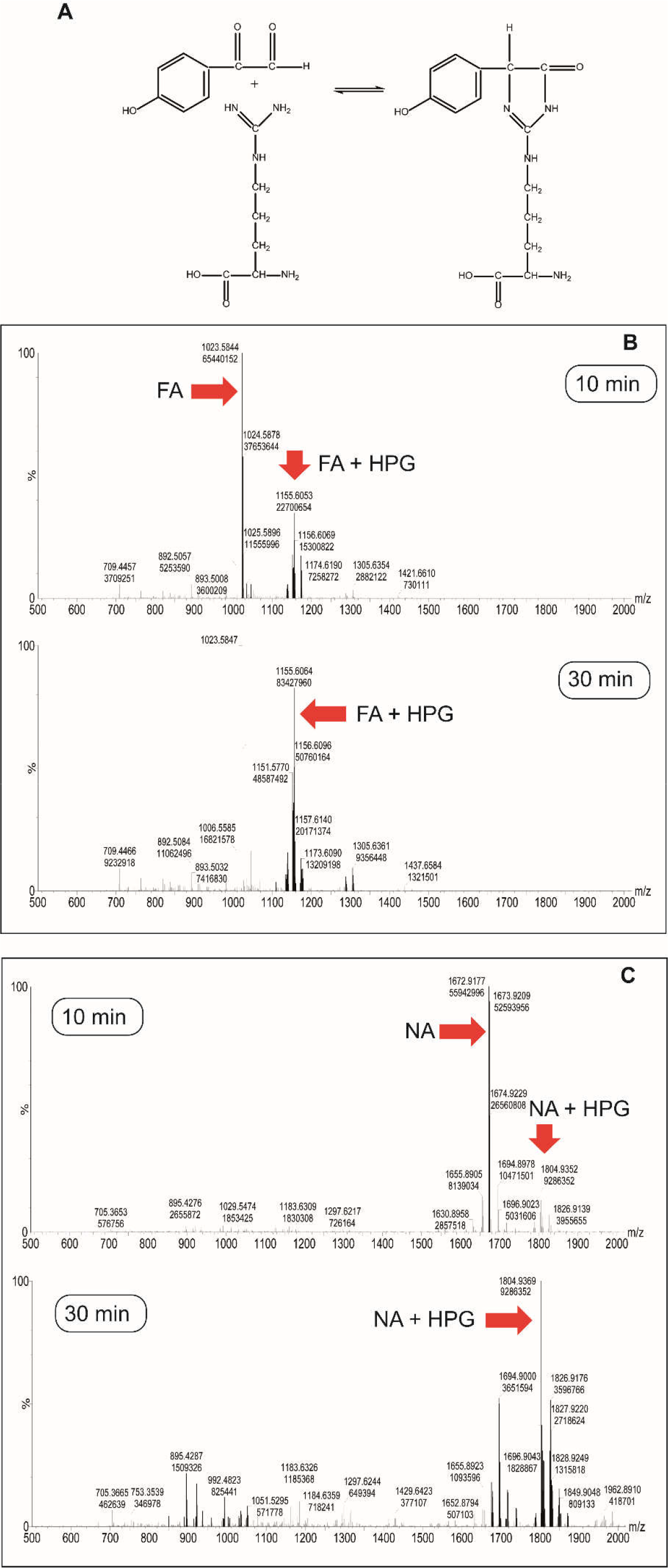
Optimization of the reaction between HPG and arginine residues on peptides. **A. The reaction of HPG and arginine.** There is a single product generated after reaction between HPG and guanidino side chain of arginine residues. **B. Mass spectra of FA reacted with HPG.** The reaction between FA and 100 mM HPG was conducted in the dark, for 10 min (Upper panel) and 30 min (Lower panel), at 25^°^C. At 10 min, the original FA with the mass of 1023.5 is observed along with and (FA+HPG) with a mass shift of 1156.6. At 30 min, only the product (FA+HPG) was observed. **C. Mass spectra of NA reacted with HPG**. The reaction between FA and 100 mM HPG was conducted in the dark, for 10 min (Upper panel) and 30 min (Lower panel), at 25^°^C. At 10 min, the original NA and along with the reaction product (NA+HPG) were detected. At 30 min, only the product (NA+HPG) was observed.

Because the model peptides used here contained a small number of arginine residues in a reasonably simple environment, it was not known whether these reaction conditions would be applicable to a protein. Thus, the next issue to address was the ability of PGO to modify all arginine side chains in a protein. Arg-C was used to digest the protein after modification and so identify the unmodified arginine residues. These experiments used FGF2, which contains eleven arginine residues, as a model protein (Fig 4).

**Figure 4:**
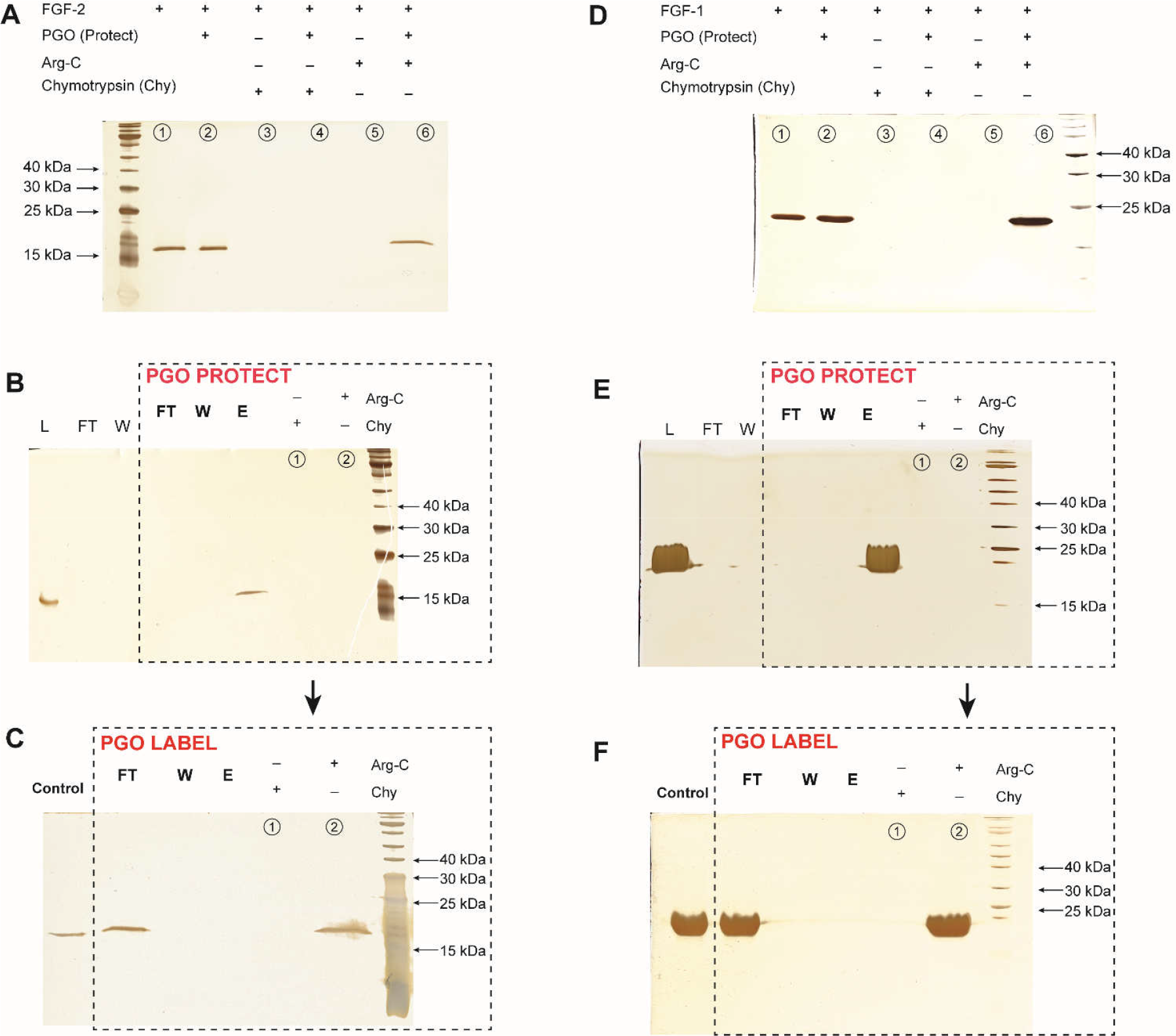
Selective labelling of arginine residues in the heparin binding sites of FGF2 and FGF1. The reaction between PGO and FGF2 or FGF1 was conducted in solution and or when bound to heparin beads in the dark, at 25^°^C for 60 min. The protein eluted from the column was further reacted with PGO in the same conditions. The products collected after reactions were incubated with chymotrypsin or Arg-C, at 37^°^C overnight. All samples were analysed by SDS-PAGE. **A. Reactions of FGF2 with PGO.** Lane 1: 1/10 of the native FGF2 used in the in solution experiment; lane 2: 1/10 of the protein after the reaction with 200 mM PGO; lane 3: products of the digestion of native FGF2 by chymotrypsin; lane 4: products of the digestion by chymotrypsin of PGO-modified FGF2; lane 5: products of the digestion by Arg-C of native FGF2; lane 6: products of the digestion by Arg-C of PGO-modified FGF2. **B. The reaction of FGF2 bound to heparin affinity column with PGO.** L, 1/10 of the protein loaded onto the column; FT, 1/10 of the flow through fraction; W, 1/10 of the fraction washed with Na-1 buffer; PGO PROTECT FT, flow through fraction after the reaction between FGF2 and PGO; PGO PROTECT W, wash fraction after the reaction with PGO; PGO PROTECT E, Eluate from the column with Na-2 buffer; PGO PROTECT lane 1: the protein in the elution was digested with chymotrypsin; PGO PROTECT lane 2: the protein in elution was digested with Arg-C. **C. Labelling of arginine residues in binding sites of FGF2.** FGF2 from the eluate (panel B, fraction PGO PROTECT E) was reacted with PGO at a final concentration of 100 mM for 60 min. The product from this the second reaction was loaded onto a heparin affinity minicolumn. L, 1/10 of the FGF2 labelled by PGO; FT, the flow through the minicolumn; W, 1/10 of the wash with Na-1 buffer; E, Elution with buffer Na-2; lane 1, the labelled FGF2 in FT fraction was digested by chymotrypsin; lane 2, the labelled FGF2 in FT fraction was digested by Arg-C. **D. Reaction of FGF1 with PGO.** Lane 1: 1/10 of the native FGF1 used in the in solution experiment; lane 2: 1/10 of the protein after the reaction with 200 mM PGO; lane 3: products of the digestion of native FGF1 by chymotrypsin; lane 4: products of the digestion by chymotrypsin of PGO-modified FGF1; lane 5: products of the digestion by Arg-C of the native FGF1; lane 6: products of the digestion by Arg-C of PGO-modified FGF1. **E. The reaction of FGF1 bound to heparin affinity column with PGO.** L, 1/10 of the protein loaded onto the column; FT, 1/10 of the flow through fraction; W, 1/10 of the fraction washed with Na-1 buffer; PGO PROTECT FT, flow through fraction after the reaction between FGF1 and PGO on the mini-column; PGO PROTECT W, wash fraction after the reaction with PGO; PGO PROTECT E, Eluate from the column with Na-2 buffer; PGO PROTECT lane 1: the protein in the elution was digested with chymotrypsin; PGO PROTECT lane 2: the protein in elution was digested with Arg-C. **F. Labelling of arginine residues in binding sites of FGF1.** FGF1 from the eluate (panel D, fraction PGO PROTECT E) was reacted with PGO at a final concentration of 200 mM for 60 min, the product of this second reaction was loaded onto a heparin affinity minicolumn. L, 1/10 of the FGF1 labelled by PGO; FT, the flow through the minicolumn; W, 1/10 of the wash with buffer Na-1; E, Elution with buffer Na-2; lane 1, the labelled FGF1 in FT fraction was digested by chymotrypsin; lane 2, the labelled FGF1 in FT fraction was digested by Arg-C.

#### The reaction of FGF2 with PGO

The reaction between FGF2 and PGO was first conducted in the absence of heparin beads. After buffer exchange, half of the protein was reacted with 200 mM PGO in the dark, at 25^0^C for 60 min (Material and Methods, step 2) and the other half was used as a control. Native FGF2 migrates as a band at round 18 kDa on SDS-PAGE (Fig 4A lane 1), but after digestion by chymotrypsin or by Arg-C no bands were apparent, demonstrating the expected cleavage of the protein by these enzymes (Fig 4A lanes 3 and 5). Reaction with PGO did not change appreciably the migration of the FGF2 (Fig 4A lane 2, PGO treated FGF2, no enzyme), and the modified protein was cleaved by chymotrypsin (Fig 4A lane 4). In contrast, PGO-modified FGF2 was not cleaved by Arg-C, since a band corresponding to FGF2 was clearly visible (Fig 4A lane 6). This suggests that all arginine residues of FGF2 were modified.

When FGF2 (Fig 4B- L) was loaded onto the heparin mini-column, no protein was observed in the flow through (Fig 4B- lane FT) and wash fractions, indicating that FGF2 bound as expected (Fig 4B- lanes FT and W). After the on-column reaction with 200 mM PGO, the excess PGO was removed (Fig 4B- lane PGO PROTECT FT) and the column was washed by Na-1 buffer (Fig 4B- lane PGO PROTECT W) before elution with Na-2 buffer (Fig 4B- lane PGO PROTECT E) (Materials and Methods, step 2). No protein was detected in the PGO PROTECT FT and PGO PROTECT W fractions, indicating that the FGF2 remained bound to heparin during the reaction. A band was observed in the elution fraction (Fig 4B-lane PGO PROTECT E), which was similar in size and amount as the loading control (Fig 4B lane L). This result indicated that protein was efficiently eluted from the column. A quarter of the protein in the eluted fraction was incubated overnight with Arg-C or chymotrypsin and the products of digestion were analysed by SDS-PAGE. In both cases, no band was apparent, indicating that the enzymes had cleaved the PGO reacted FGF2 (Fig 4B PGO PROTECT lanes 1 and 2). Thus, when FGF2 is reacted with PGO in solution, there is a complete modification of arginine residues, but when the reaction is performed on FGF2 bound to heparin, the modification is incomplete. This suggests that only arginine residues exposed to solvent in heparin bound FGF2 were able to react with PGO.

The remaining half of the eluted protein (~20 μL) was diluted with Na-1 buffer until the final concentration of NaCl was less than 0.2 M and then reacted in solution with PGO at 100 mM final concentration. Half of this product was applied onto a mini heparin affinity column. The flow-through fraction contained a band of almost identical intensity to the reaction product (Fig 4C, lane FT) and there was no protein detected in the wash (Fig 4C, lane W) and eluted fractions (Fig 4C, lane E) indicating that the FGF2 no longer bound to heparin. Thus, after the second reaction (equivalent to the labelling step in the original lysine selective protect and label [11]) the arginine residues in HBSs were blocked by PGO. The FGF2 reacted with PGO a second time and recovered from the flow through fraction was also probed with proteases. While chymotrypsin digested the FGF2 (Fig 4C lane 1), Arg-C was unable to do so, since there was a band (Fig 4C lane 2) at the same size as the initial reaction product (Fig 4D -L) and labelled FGF2 (Fig 4C - FT). Hence, the arginine residues protected after the initial on-column reaction with PGO were successfully blocked in the second reaction of this FGF2 with PGO in solution.

These data suggest that under the reaction conditions used PGO successfully modified the side chains of all 11 arginine residues of FGF2, and that following PGO modification of those arginine residues that remain exposed when FGF2 binds to heparin, the remaining arginine residues on the eluted protein may also be successfully modified.

#### The reaction of FGF1 with PGO

The same series of experiments were repeated with FGF1. In the absence of heparin beads, after buffer exchange, half of the protein was reacted with 200 mM PGO in the dark, at room temperature for 60 min (Material and Methods, step 2) and the other half was used as a control. The expected cleavage of the native FGF1 (Fig 4D lane 1) by chymotrypsin or by Arg-C was observed, evidenced as no detectable bands (Fig 4D lanes 3 and 5). Whereas the PGO-modified FGF1 was cleaved by chymotrypsin (Fig 4D lane 4), it was not cleaved by Arg-C, since a band corresponding to FGF1 was clearly visible in this case (Fig 4D lane 6). This suggests that all arginine residues of FGF1 were modified.

When FGF1 (Fig 4E- L) was loaded onto the heparin mini-column, no protein was detected in the flow through (Fig 4E lane - FT) and wash fractions,nor when PGO was applied (Fig 4E - PGO PROTECT FT and PGO PROTECT W lanes), indicating that the FGF1 remained bound to heparin during the reaction. A band was observed in the elution fraction (Fig 4E- lane PGO PROTECT E) lane, which was similar in size and amount as the loading control (Fig 4E L lane). This result indicated that protein was efficiently eluted from the column. When a quarter of the PGO reacted FGF1 was incubated overnight with Arg-C or chymotrypsin, no band was apparent (Fig 4E PGO PROTECT lanes 1 and 2), demonstrating that there remained unreacted arginine residues. After a second reaction of eluted protein with PGO at 200 mM final concentration, chymotrypsin digested the FGF1 (Fig 4F lane 1), but Arg-C was unable to do so, since there was a band (Fig 4F lane 2) at the same size as the initial reaction product (Fig 4F, lane L) and FGF1 reacted with PGO in solution (Fig 4F, FT lane). Moreover, this eluted FGF1 subjected to a second reaction in solution with PGO failed to bind to heparin column (Fig 4F, lane FT). Hence, the arginine residues engaged with heparin in the initial on-column reaction with PGO were successfully reacted in the second reaction of this FGF1 with PGO in solution.

#### Protect and Label strategy for the identification of arginine residues in FGF1 and FGF2 that bind heparin

There is a large body of published structural, biophysical and biological data relating to the interactions of FGF1 and FGF2 with heparan sulfate and its experiment proxal heparin. FGF1 and FGF2 share a high level of similarity in structure and sequence, though they possess very different isoelectric points, 6.52 and 11.18, respectively. FGF1 and FGF2 were loaded on to AF-heparin mini columns and reacted *in situ* with 200 mM PGO (Fig 1) (Material and Methods, step 1). The eluted proteins were then reacted for 60 min with 100 mM HPG in the dark (Material and Methods, step 3) and processed for mass spectrometry (Material and Methods, step 4). In parallel, FGF1 and FGF2 were reacted with 200 mM PGO in the absence of heparin beads (Material and Methods, step 2). The native and modified proteins were then cleaved by chymotrypsin (Material and Methods, step 5).

Peptides produced from digestions were predicted using two protein identification and analysis tools, Protein Prospector (v 5.19.1 developed by the USCF Mass spectrometry Facility) and Peptide Mass (ExPASy). In both cases, these provided parameters included the mass range from 500 Da to 4000 Da, maximum missed cleavages 5, monoisotopic only and the enzyme used. For Protein Prospector, the oxidation of methionine was considered as the variable modification, whereas it was not included by Peptide Mass (ExPASy).

To identify the reaction products of the arginine side chain and PGO, their structures and masses were evaluated (Table 2). Depending on the stoichiometry of the reactions and the loss of water, there were four possible products: 1:1 (1PGO: 1arginine), (Product 1- Fig 2A); 1:1 water-condensed, (Product 2- Fig 2A) and 2:1 product water-condensed, (Product 3, 4- Fig 2A). Product 3 and 4 had different structures, but they led to the same mass shift for the reacted arginine residue. The corresponding mass shifts resulting from these products are provided in Table 2. The 250 kDa product (Supplementary Fig 2C) was not considered, as neither of the two FGFs possess a terminal arginine. To identify arginine residues protected by heparin binding, a second reagent was required, that would react similarly with arginine side chains, but yield a product with a different mass. HPG was chosen for this purpose, and this yields just a single, 1:1 product (Table 2, Fig 3).

**Table 2:** Products of PGO/HPG and arginine side chains with corresponding mass shift. The summary of the products between PGO/HPG and arginine side chains. The first PGO reacts with the guanidino group to produce1:1 product (Fig 2A, product [1]), which could then reversibly release a hydroxyl group from glyoxal, resulting in the 1:1 product water-condensed product (Fig 2A, product [2]). Following is the reaction of a second PGO molecule (if sufficient reactant) onto either a nitrogen of the guanidino group of arginine (Fig 2A, products [3]) or the carbonyl group of the first PGO (Fig 2A, products [4]) to form the final 2:1 product water-condensed product.

| # of product | Product | Mass shift |
| --- | --- | --- |
| Figure 2A 1 | 1:1 product | 134.04 |
| Figure 2A 2 | 1:1 product water-condensed | 116.02 |
| Figure 2A 3 | 2:1 product | 252.04 |
| Figure 2A 4 | 2:1 product water-condensed | 232.04 |
| Figure 3A | HPG | 132.02 |

Peptides from the native and modified proteins produced by enzyme cleavage were analysed by MALDI-Q-TOF mass spectrometry. Following was the analysis of the mass spectra using scripts *PGO-HPG mass predictor* and *Matchmaker*.

#### Identification of arginine residues in FGF2 involved in binding heparin

FGF2 has eleven arginine residues in the sequence, R^31^, 42, 48, 53, 69, 81, 106, 116, 118 and 129. The peptides generated from native FGF2 by cleavage with chymotrypsin were predicted by Peptide Mass and Prospector, then filtered with the script PGO-HPG mass predictor to remove peptides without arginine residues. Prospector predicted 158 peptides and there were 126 peptides predicted by Peptide Mass with 100% sequence coverage, demonstrating that all arginine residues could be analysed.

##### Protection of arginine residues on FGF2 by PGO in solution

To understand the accessibility of the reagent to arginine, FGF2 was reacted with 200 mM PGO for 60 min in 0.2 M NaHCO_3_, pH 9.5. Peptides of the modified FGF2 were generated by cleavage with chymotrypsin and analysed by mass spectrometry. The modification on each arginine residue was identified by *PGO-HPG mass predictor* and *Matchmaker*. The resulting peptides with information about modification, sequence,and final m/z are presented in Supplementary Tables 1 and 2 and the spectra are in Supplementary Fig 3.

Among peptides predicted by Prospector, nine of them contained the reacted arginine residues (Supplementary Table 1, Supplementary Fig 3). PGO product 1 (Fig 2A) was observed in peptide 2 with R^129^, peptides 4, 5 and 8 with R^116^ and R^118^, peptide 7 with R^69^ and R^81^. R^90^ of peptide 3 reacted with PGO to generate product 2 (Fig 2A). Peptide 1 with R^31^, peptide 6 with R^106^ showed a mass shift corresponding to products 3, 4 (Fig 2A). A mixture of products was found in peptide 9 with R^42^, 48 and 53, as a combination of products 1 and 3. It was noticed that only two out of three arginines on peptide 9 reacted with PGO, indicating that one arginine was apparently not accessible to PGO in this context. Hence, the products of ten arginine residues with PGO were identified, though a lack of reactivity of one arginine to the reagents was also observed.

Using the predicted peptides from ExPASy, the number of peptide with modified arginine residues was six, which completely overlapped with the list generated by Prospector (Supplementary table 2).

##### Selective labelling of arginine residues on FGF2 by PGO and HPG on heparin mini column

PGO-HPG mass predictor and Matchmaker scripts were used to identify the peptides containing the arginine residues that are labelled by HPG and therefore engaged with heparin. The resulting peptides with information about modification, sequence, and final m/z are presented in Tables 3 and 4 and the spectra are in Supplementary Figs S4 and S5).

**Table 3:** FGF-2 peptide analysis based on prediction by Prospector. The reference number of the peptide is followed by the predicted m/z and with the sequence of the peptide following cleavage of native FGF-2 by chymotrypsin. The two columns under “Native FGF-2” present the predicted m/z and sequences of peptides after chymotrypsin digestion of FGF-2.The first two columns under “FGF-2 protect and label (P&L)” present the observed and predicted m/z for FGF-2 after modifications of arginine residues. The third column is the assignment of the peptide to one of the three HBSs of FGF2[11,13]. The final columns indicates the modification occurring on the arginine residues of the peptides.

|  | Native FGF-2 |  | FGF-2 protect and label (P&L)<sup>1</sup> |  |  |  |
| --- | --- | --- | --- | --- | --- | --- |
|  | m/z theoretical | Sequence | m/z observed | m/z theoretical | Published HBS<sup>2</sup>. | Modifications |
| **1** | 996.54 | <sup>116</sup>**RSRKY**TSW<sup>123</sup> | 1144.62 | 1144.56 | HBS2 (R<sup>116</sup> and R<sup>118</sup>) | 1 R + 132 (**HPG**) and oxidation |
| **2** | 1032.55 | <sup>83</sup>**AMKEDGR**LL<sup>92</sup> | 1180.59 | 1180.59 | HBS2 (R<sup>90</sup>) | 1 R + 132 (**HPG**) and oxidation |
| **3** | 1032.55 | <sup>82</sup>**LAMKEDGR**L<sup>91</sup> | 1180.59 | 1180.59 | HBS2 (R<sup>90</sup>) | 1 R + 132 (**HPG**) and oxidation |
| **4** | 1318.61 | <sup>105</sup>FFERLESNNY<sup>115</sup> | 1434.63 | 1434.61 |  | 1 R + 116(**PGO**) |
| **5** | 1374.69 | <sup>113</sup>NTY**RSRKY**TW<sup>123</sup> | 1738.82 | 1738.74 | HBS2 (R<sup>118</sup>) | 2 R + 132 (**HPG**) + 232 |
| **6** | 1439.84 | <sup>124</sup>YVAL**KRTGQYKL**<sup>135</sup> | 1587.85 | 1587.86 | HBS1 (R<sup>129</sup>) | 1 R + 132 (**HPG**) and oxidation |
| **7** | 1525.79 | <sup>27</sup>**KDPKRLYCK**NGGF<sup>40</sup> | 1673.79 | 1673.82 | HBS3 (R<sup>31</sup>) | 1 R + 132 (**HPG**) and oxidation |
| **8** | 1979.01 | <sup>65</sup>QAEERG**VVS****IKG**VCANRY<sup>82</sup> | 2211.15 | 2211.08 | HBS2 | 1 R + 232(**PGO**) |
| **9** | 2220.16 | <sup>63</sup>QLQAEERG**VVS****IKG**VCANRY<sup>82</sup> | 2353.17 | 2353.15 | HBS2 (R<sup>69</sup> and R<sup>81</sup>) | R + 133 (**PGO**) |
| **10** | 2537.41 | <sup>41</sup>LRIHPDGRVDGVREKSDPHIKL<sup>62</sup> | 2769.45 | 2769.45 |  | 2 R + 116 x 2(**PGO x2**) |

<sup>1</sup> P&L: FGF-2 reacted in solution with PGO when bound to a heparin affinity column (protection step) and then, following elution, reaction of any protected arginine residues with HPG (labelling step).
<sup>2</sup> The bolded amino acids are a part of the known HBSs which were noted in column 3<sup>rd</sup> of FGF-2 on column.

**Table 4:** FGF-2 MS analysis based on prediction by Peptide Mass (ExPASy). The reference number of the peptide is followed by the predicted m/z and with the sequence of the peptide following cleavage of native FGF-2 by chymotrypsin. The two columns under “Native FGF-2” present the predicted m/z and sequences of peptides after chymotrypsin digestion of FGF-2.The first two columns under “FGF-2 protect and label (P&L)” present the observed and predicted m/z for FGF-2 after modifications of arginine residues. The third column is the assignment of the peptide to one of the three HBSs of FGF2[11,13]. The final columns indicates the modification occurring on the arginine residues of the peptides.

|  | Native FGF2 |  | FGF-2 (P&L)<sup>3</sup> |  |  |  |
| --- | --- | --- | --- | --- | --- | --- |
|  | m/z theoretical | Sequence | m/z observed | m/z theoretical | Published HBS<sup>4</sup> | Modifications |
| **1** | 996.54 | <sup>116</sup>**RSRKY**TSW<sup>123</sup> | 1144.63 | 1144.56 | HBS2 (R<sup>116</sup> and R<sup>118</sup>) | 1 R + 132 (**HPG**) and oxidation |
| **2** | 1032.55 | <sup>82</sup>**LAMKEDGRL**<sup>91</sup> | 1180.59 | 1180.58 | HBS2 (R<sup>90</sup>) | 1 R + 132 (**HPG**) and oxidation |
| **3** | 1032.55 | <sup>83</sup>**AMKEDGRLL**<sup>92</sup> | 1180.59 | 1180.58 | HBS2 (R<sup>90</sup>) | 1 R + 132 (**HPG**) and oxidation |
| **4** | 1198.66 | <sup>124</sup>**YVALKRTGQY**<sup>134</sup> | 1362.61 | 1362.67 | HBS1 (R<sup>129</sup>) | 1 R + 132 (**HPG**) and oxidation |
| **5** | 1276.77 | <sup>125</sup>**VALKRTGQYKL**<sup>135</sup> | 1408.71 | 1408.77 | HBS1 (R<sup>129</sup>) | 1 R + 132 (**HPG**) |
| **6** | 1318.61 | <sup>105</sup>**FFERLESNNY**<sup>115</sup> | 1434.63 | 1434.61 |  | 1 R + 116 (**PGO**) |
| **7** | 1374.69 | <sup>113</sup>**NTYRSRKYTW**<sup>123</sup> | 1738.82 | 1738.74 | HBS2 | 2 R + 132 (**HPG**) + 232 |
| **8** | 1439.84 | <sup>124</sup>**YVALKRTGQYKL**<sup>135</sup> | 1587.85 | 1587.86 | HBS1 (R<sup>129</sup>) | 1 R + 132 (**HPG**) |
| **9** | 1525.79 | <sup>27</sup>**KDPKRLYCKNGGF**<sup>40</sup> | 1673.79 | 1673.82 | HBS3 (R<sup>31</sup>) | 1 R + 132 (**HPG**) |
| **10** | 1979.01 | <sup>65</sup>**QAEERGVVSIKGV**CANRY<sup>82</sup> | 2211.15 | 2211.08 | HBS2 | 1 R + 232(**PGO**) |
| **11** | 2220.15 | <sup>63</sup>**QLQAEERGVVSIKGV**CANRY<sup>82</sup> | 2369.17 | 2369.15 | HBS2 (R<sup>69</sup> and R<sup>81</sup>) | 2 R + 133(**PGO**) |
| **12** | 2537.41 | <sup>41</sup>**LRIHPDGRVDGVREKSDPHIKL**<sup>62</sup> | 2769.27 | 2769.26 |  | 2 R + 116 x 2(**PGOx2**) |

<sup>3</sup> P&L: FGF-2 reacted in solution with PGO when bound to a heparin affinity column (protection step) and then, following elution, reaction of any protected arginine residues with HPG (labelling step).
<sup>4</sup> The bolded amino acids are a part of the known HBSs which were noted in column 3<sup>rd</sup> of FGF-2 on column.

After the protection step on the heparin affinity column with PGO and labelling with HPG ten peptides with modified ariginines were identified (Table 3) (Supplementary Fig 4). Of these, four peptides (peptides 4, 8, 9 and 10) contained PGO protected arginine residues only; peptides 1, 2, 3, 6 and 7 had a mass shift corresponding to the product of HPG (132.02 Da) whereas, in peptide 5, a mixture of PGO and HPG products was observed (Table 3). The first arginine on the sequence is R^31^, situated in the disordered N-terminal region adjacent to beta strand I. Peptide 7 containing R^31^ had a mass shift corresponding to HPG (Table 3), indicating that this arginine engages heparin. From previously defined HBSs of FGF2, R^31^ is part of HBS-3 which locates to the N-terminus of the beta trefoil FGF (Fig 7).

Following on in the sequence are R^42^, R^48^, and R^53^, which were detected on peptide 10. The mass shift of this peptide corresponded to two 1:1 condensed products between arginine and PGO. Hence, only two of the three arginines were modified by PGO, whereas one remained unreactive to PGO/HPG(Table 3). This peptide was also observed with just two modifications by PGO when the reaction was performed with FGF2 in solution in the absence of heparin (Supplementary table 1, peptide 9). The presence of an unlabelled arginine is surprising, since Arg-C could not cleave FGF2 after reaction with PGO and HPG (Fig 4A-C). The question of which arginine was not accessible to PGO, HPG or trypsin is addressed later.

Next, Arg^69^ and Arg^81^ were protected by PGO, as observed in two sister peptides 8 and 9. They cover the sequence from the loop between strand IV-strand V to a loop between strand V-strand VI, in which K^75^ has been defined as part of HBS-2. Peptide 9 is two amino acids longer than its sister, but their reaction products are distinct. While peptide 8 showed a single mass shift corresponding to the 2:1 condensed product, the 1:1 product was observed in peptide 9, but only one of it's two arginine reacted with PGO and the other failed to react with PGO or HPG. The MALDI-TOF data are not conclusive as to which arginine was modified in each case and to whether PGO reacted with one arginine in both cases or it reacted with different arginine residues. Interestingly, when FGF2 reacted with PGO in-solution, Arg^69^ and Arg^81^ were both modified by PGO generating two 1:1 products (Supplementary table 1, peptide 7). These differences between reaction products may be due tothe effect of either higher concentrations of electrolytes in the reaction of the eluted protein with HPG or to a long-lasting effect on protein conformation of heparin binding.

Two sister peptides, 2 and 3, containing R^90^ from strand VI/strand VII loop (Fig 7) were labelled by HPG (Table 3), indicating that this residue was involved in the binding of FGF2 to heparin. R^90^ has not been shown in previous publications to be part of a HBSs (Fig 7). Peptide 4 covers the sequence from F^105^ to Y^115^, containing Arg^108^ in strand VIII (Fig 7) which formed the 1:1 condensed product with PGO, resulting in the mass shift of 116 Da.

Peptides 1 and 5 contain two arginine residues, R^116^ and R^118^ in strand IX, assigned previously though sequence alignment to HBS-2 (Fig 7) [13]. Peptide 1 showed a mass shift corresponding to one HPG, indicating that one of these two adjacent arginines engaged with heparin (Table 3). In peptide 5 there were two modifications, one from reaction with HPG, one from reaction with PGO (Table 3). In solution, both R^116^ and R^118^ were modified by PGO generating two 1:1 water condensed products (peptide 4 and peptide 5, Supplementary table 1). To answer the question of which of the arginine residues was bound to heparin, a protection reaction with PGO on a heparin mini-column was followed by sequential digestion for 5 hours with chymotrypsin then overnight with Arg-C. The peptides were analysed by MALDI_TOF MS. As result, R^118^ of peptide 5 (Table 3) was cut by Arg-C, resulting in peptide ^113^NTYRSR^118^ with a mass before modification of 795.4 and a mass after modification of R^116^ by PGO of 1027.13 (Supplementary Fig 6). This observation demonstrated that R^118^ binds to heparin whereas R^116^ does not. The different behaviour of peptides 1 and 5 may be due to the N-terminal location of the arginine residue causing the loss of a PGO in some instances.

The last arginine on the sequence, R^129^ in strand X/strand XI loop (Fig 7) showed a mass shift of 132.02 corresponding to the modification by HPG (peptide 6, Table 3). This agrees with its prior assignment to HBS-1[11][13].

Using Peptide Mass (ExPASY) to predict peptides generated from native FGF2 by cleavage with chymotrypsin, Matchmaker found 12 modified peptides, summarized in Table 4 (Supplementary Fig 4). Ten of them overlap with the list generated by Prospector prediction. The extra two peptides were sisters that showed a mass shift corresponding to HPG on R^129^ of the canonical HBS-1 (Table 4).

#### Locations of modified arginine residues in the FGF2 structure

The canonical HBS of FGF2, HBS-1, has been characterized by many different methods, including X-ray crystallography, NMR spectroscopy, and site-directed mutagenesis, while evidence for its two secondary binding sites, HBS-2 and HBS-3, has also been acquired by independent approaches [11,17,18,20,36,37]. The primary binding site (HBS1) consists of Lys^35^ and Asn^36^ and the group of 17 amino acids from 128-144 (Fig 7) [17]. In addition, HBS-3 was identified towards the N-terminus, which consists of 5 amino acids in the region 25-30, and HBS-2 is formed by Lys^75^ and Gly^76^, ^82^LMAK^86^ and Lys^119^ [11,38].

##### HBS-1, Arg129 and Arg90

One arginine in HBS-1, R^129^, was labelled by HPG (peptide 6_Table 3; peptides 4, 5 and 8_Table 4). HBS-1 is an extremely basic surface on FGF2 (Fig 5A) formed by Lys35, Asn36 on strand I/strand II loop and residues in strand X/strand XI loop, strand XI and strand XI/strand XII loop (Fig 7). The double mutation R129A/K143A dramatically reduced the binding of the protein to a heparin affinity column [17]. In addition, R^129^ is highly conserved inthe FGF family and it aligns to K^128^ of FGF1, which engages heparin, as shown previously[13].

**Figure 5:**
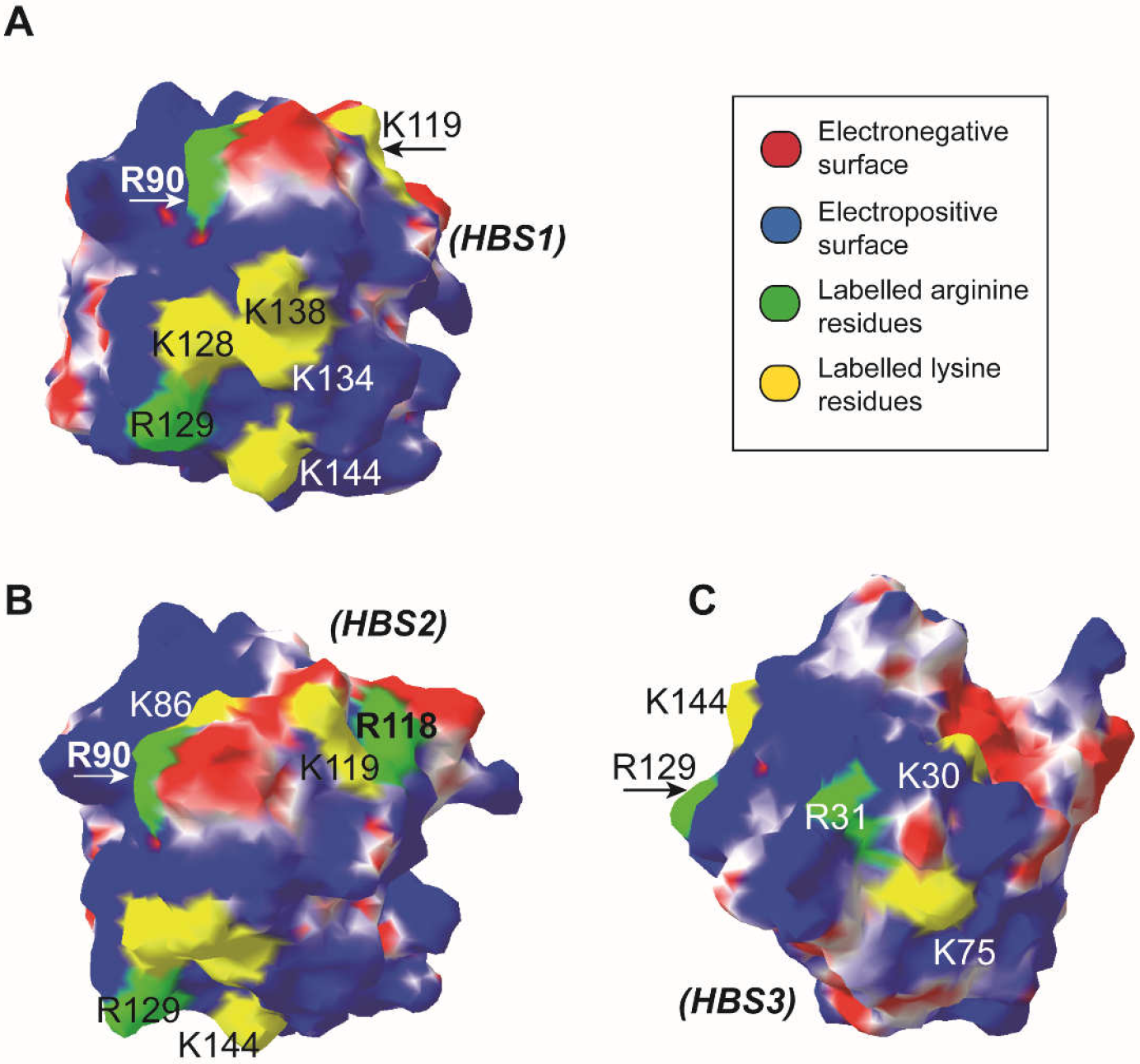
Labelled arginine and lysine residues in heparin-binding sites of FGF2. The FGF2 structure (PBD code **1bfg** [44]) is shown as a surface.The electrostatic potential of FGF2 and FGF1 was computed using Poisson-Boltzmann algorithm by Swiss-PDBView with the positively charged areas coloured *blue* and the negatively charged areas coloured *red*. Labelled lysine residues [11,13] are presented in *yellow*. Labelled arginine residues (Tables 3, 4) are coloured *green*. This colour scheme for electrostatic potential is used throughout the paper including Figs 5,6. **A. Basic residues in HBS1.** R^129^ of the canonical HBS-1. Other basic residues in HBS1 include K^128^, K^134^, K^138^, K^144^ [11,13]. R^90^ and K^86^ which are now assigned to be part of HBS1 by this work are shown. **B**. **Basic residues in HBS2.** Location of R^118^ and K^119^ are shown on the surface in relative to the rest of HBS-2. All the basic residues in HBS1 can also be seen from this viewpoint as well. **C**. **Basic residues in HBS3.** Location of K^30^, R^31^ and K^75^.

The location of R^90^ on the surface of FGF2 has suggested that this arginine is a part of HBS-1 (Fig 5A). Peptides 2 and 3 (Table 3 and Table 4, respectively) demonstrated that R^90^ reacted with HPG and hence is bound to heparin. The involvement of R^90^ in heparin binding has been suggested by a docking model to be an indirect interaction through an intervening water molecule to the GlcNSO3^-^ group of glucosamine 5 (GlcNS, -5) at the reducing end of the sugar ligand [17][18].

##### HBS-2, Arg118

HBS-2 comprises amino acids scattered in sequence, Lys^75^, K^86^ and ^116^RSRK^119^. Sequence alignment, following the selective labelling of lysine residues, suggested that R^118^ and R^116^ were part of HBS2 [30]. However, the selective labelling of arginine residues demonstrates R^118^ but not R^116^ is bound to heparin. ^118^RK^119^ is separated from HBS1 by an acidic boundary (Fig 5B), and hence likely to constitute a distinct binding site.

##### Re-assignment of K^86^ to HBS1

The selective labelling of lysine has identified K^86^ as part of the secondary HBS2 [13]. However, on the surface, K^86^ is adjacent to R^90^ of HBS1 and situated on a positively charged region (Fig 5B). These observations prompted us to propose the re-assignment of K^86^ to HBS1 rather than HBS2.

##### HBS-3, Arg31

HBS-3 is located N terminal to strand beta I (Fig 7). The lysine selective labelling of FGF2 found K^30^ to be biotinylated and K^27^ to be acetylated, implying the involvement of K^30^ in heparin engagement [13]. R^31^ reacted with HPG (Peptide 7 – Table 3, Peptide 9 – Table 4) so can be considered to be part of HBS3.

##### Re-assignment of K^75^ to HBS3

Interestingly, K^75^, which was considered as part of HBS2 by the selective labelling of lysine [13], is distant from ^118^RK^119^ of HBS2. On the other hand, it is close to K^30^ and R^31^ of HBS3 (Fig 5C) in a continuous positively charged area of the protein’s surface. Hence, we propose that K^75^ of FGF2 is part of HBS3, not HBS2.

##### HBS3 has two mutually exclusive partners: HS and FGFR

The structure of a co-crystal of FGF2 and FGFR1c (1CVS) [39] indicated that N-terminal segment of FGF2 interacts with the third immunoglobin loop of FGFR1. In particular, K^31^ of HBS3 forms a hydrogen bond with the side chain of Gln-284 and Asp-282 of FGFR1 [39]. In addition, sequence alignment data demonstrated that K^30^ and R^31^ were aligned to R^84^ and R^85^ of FGF4, respectively, which have been shown to interact with FGFR1 [40]. These observations imply that HS and FGFR are mutually exclusive binding partners of HBS3, and these amino acids may switch partners during the formation of the receptor signalling complex.

##### Arg42 of FGF2 is trapped in an intramolecular network of hydrogen bonds

R^42^ of FGF2 (peptide 3 – Table 4) was not modified by PGO in the in solution context or on the heparin affinity mini column and was not a site for Arg-C cleavage. This implies that this residue may have intramolecular interactions, which are sufficient to prevent significant interactions with solvent (Supplementary Fig 7). The stick structure of FGF2 illustrates that the side chain of R^42^ is 0.29 nm and 0.38 nm from the side chain of D^50^; 0.29 nm and 0.38 nm to that of D^57^ and 0.34 nm and 0.38 nm to that of V^52^ (Supplementary Fig 7). These measurements suggested that the guanidino group of R^42^ is engaged in an intramolecular hydrogen bond network with the side chains of D^50^ and D^57^, the invariant residues in the FGF family, with the side chain of V^49^ forming a hydrophobic environment for the aliphatic portion of the arginine side chain. These interactions hold the side chain of R^42^ and so prevent the access of Arg-C and PGO/HPG. This intramolecular network of R^42^, D^50^, V^52^,and D^57^ may serve to restrict the conformational freedom of the four antiparallel β strands I, II, III and IV.

#### Identification of arginine residues on FGF1 engaging heparin by Protect and Label strategy

FGF1, originally called acidic fibroblast growth factor due to its isoelectric point (pI) 6.52, has six arginine residues of which, only R^134^ and R^137^ located in HBS1 have been previously identified as interacting with heparin. The other arginine residues are R^39^, 50, 52 and 103. Solvent-exposed arginine residues in heparin-bound FGF1 were protected with PGO and following elution of the FGF1, any arginine side chains engaged with the polysaccharide were labelled with HPG. The resulting peptides with information about modification, sequence, and final m/z are presented in Tables 5 and 6 and the spectra are in Supplementary Figs 8 and 9).

**Table 5:**
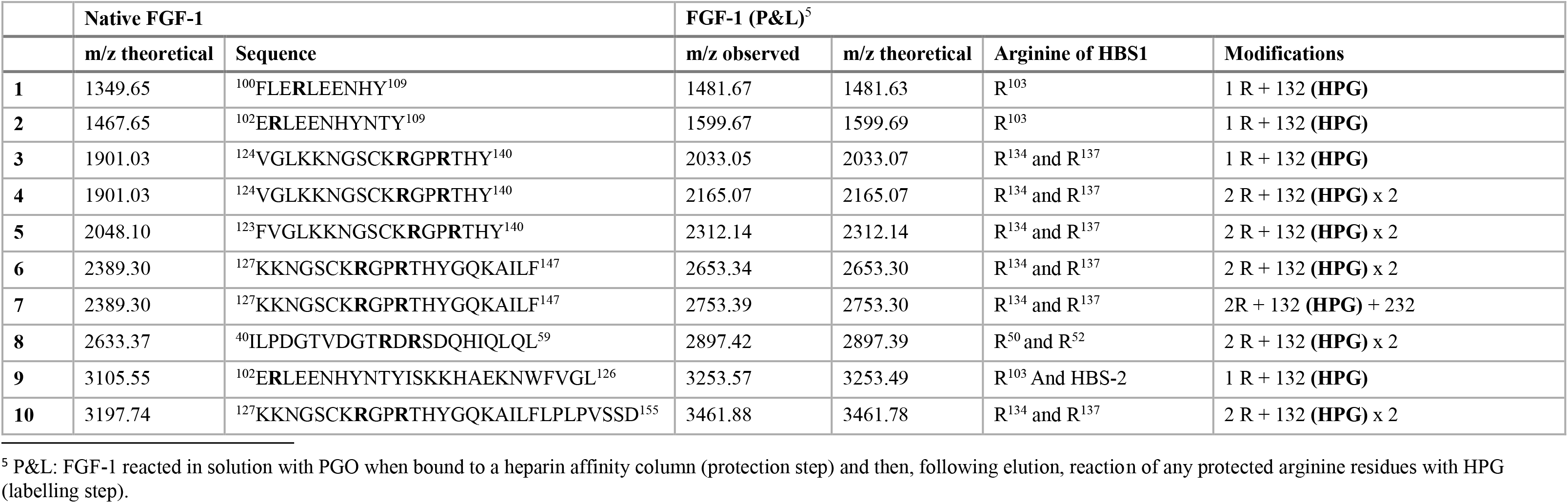
FGF-1 MS analysis based on prediction by Prospector. The reference number of the peptide is followed by the predicted m/z and with the sequence of the peptide following cleavage of native FGF-1 by chymotrypsin. The two columns under “Native FGF-1” present the predicted m/z and sequences of peptides after chymotrypsin digestion of FGF-1.The first two columns under “FGF-1 protect and label (P&L)” present the observed and predicted m/z for FGF-1 after modifications of arginine residues. The third column is the location of arginine residues of HBS-1 of FGF-1. The final columns indicates the modification occurring on the arginine residues of the peptides.

**Table 6:**
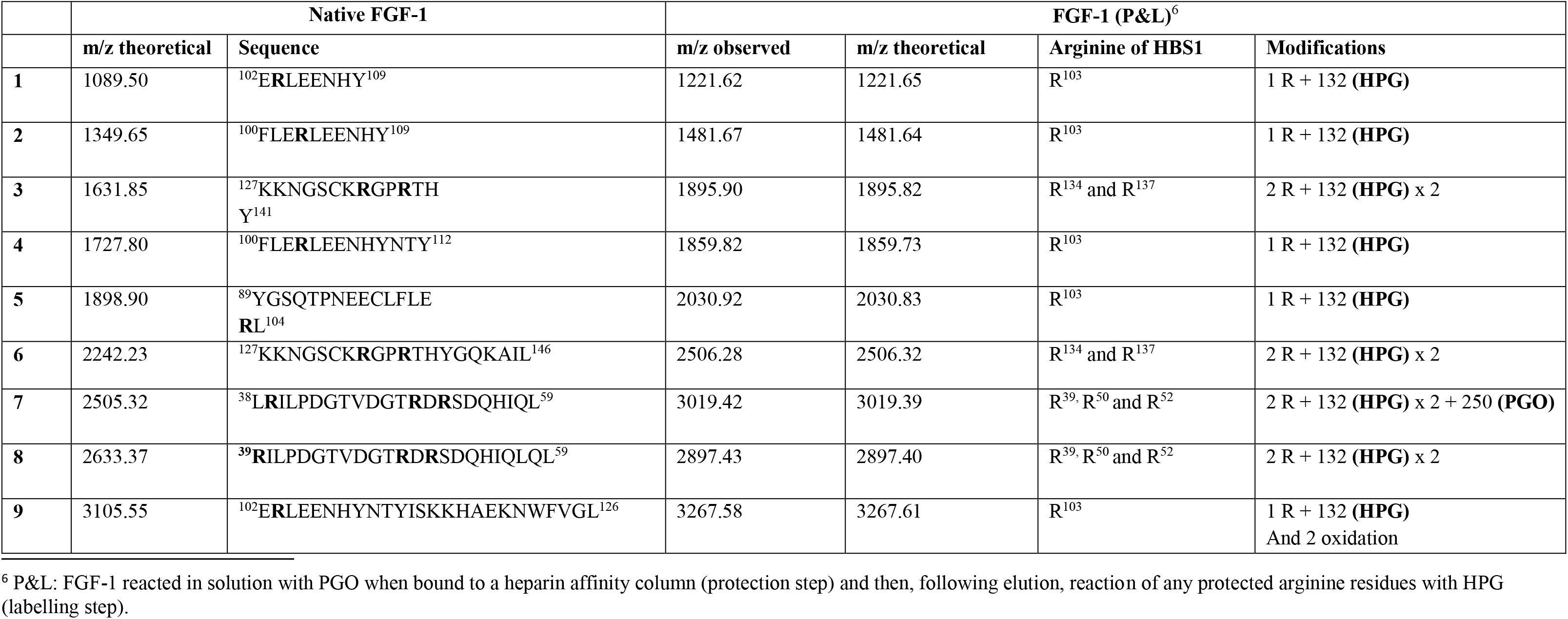
FGF-1 MS analysis based on prediction by Peptide Mass. The reference number of the peptide is followed by the predicted m/z and with the sequence of the peptide following cleavage of native FGF-1 by chymotrypsin. The two columns under “Native FGF-1” present the predicted m/z and sequences of peptides after chymotrypsin digestion of FGF-1.The first two columns under “FGF-1 protect and label (P&L)” present the observed and predicted m/z for FGF-1 after modifications of arginine residues. The third column is the location of arginine residues of HBS-1 of FGF-1. The final columns indicates the modification occurring on the arginine residues of the peptides.

##### Prospector

Prospector predicted 148 peptides after cleavage of FGF1 by chymotrypsin with 100% sequence coverage, indicating that all six arginine residues would be included in the analysis. However, analysis of the data with Prospector did not identify a peptide containing R^39^ (Table 5) indicating that coverage was incomplete. Subsequently, the filter and analysis identified ten peptides with HPG-labelled arginines (Table 5) (Supplementary Fig 8).

R^50^ and R^52^ from beta strand III/beta strand IV loop (Fig 7) were found in peptide 8 (Table 5) and this showed a mass shift of two HPG products, indicating that both arginines were bound to heparin. R^103^ from beta strand VIII (Fig 7) reacted with HPG as observed in peptides 1, 2, 9 (Table 5), indicating the binding of this arginine to heparin.

The engagement of R^134^ and R^137^ in the HBS1 of FGF1 to heparin was demonstrated by HPG modification and a mass shift of two HPG products were found in peptides 4, 5, 6 and 10 (Table 5). However, peptide 3, which also contains R^134^ and R^137^, only showed the mass shift of one HPG (Table 5, Supplementary Fig 13), meaning that one arginine had not reacted.Peptide 3 is identical to peptide 4 and overlaps with peptides 6 and 10. The mass shift of peptide 7 (Table 5, Supplementary Fig 8) is a combination of one HPG product and one 2:1 condensed product of reaction with PGO. Together these data suggest that while R^134^ and R^137^ are engaged to heparin, one of them may dissociate in the time of the protection step and so react with PGO, and moreover, may engage in intramolecular bonds rendering it resistant to reaction.

##### Peptide Mass (ExPASY)

In the case of FGF2, Prospector and ExPASY demonstrated a high level of overlap in terms of peptides containing modified arginine residues but, their predications are more diverse for FGF1. Nine peptides with modified arginines were identified when Peptide Mass (ExPASY) was used as the starting point for the analyses (Table 6, Supplementary Fig 9), but only two of them appeared in the list generated by Prospector, peptides 2 and 8.

R^39^, R^50^, and R^52^ were in peptides 7 and 8 (Table 6). Three products are observed in peptide 7, in which, only two products of arginine and HPG were detected, the other is a 2:1 product of PGO and arginine. In the case of peptide 8, the mass shift was attributed to two products of arginine with HPG, implying a lack of long-lasting reaction between one arginine with PGO or HPG. With the additional evidence from peptide 8 (Table 3) generated byProspector, it could be concluded that HPG reacted with R^50^ and R^52^.

R^103^ from beta strand VIII was labelled with HPG, resulting in a mass shift of HPG in five peptides, 1, 2, 4, 5 and 9 (Table 6), demonstrating its involvement in heparin binding. R^134^ and R^137^ of HBS-1 were labelled by HPG as observed in peptides 3 and 6 (Table 6).

#### Locations of modified arginine residues in the FGF1 structure

Previous work demonstrated that FGF1 has three regions on its surface that engage heparin: the canonical HBS-1 and the secondary binding sites, HBS-2 and HBS-3[13,41,42]. The core of the canonical heparin binding site of FGF1 is almost aligned to that in FGF2 (Fig 7), and extends from beta strand IX/beta strand X loop to beta strand XI/beta strand XII loop (Fig 7) as evidenced by X-ray crystallography and NMR spectroscopy[13,43–45]. Lysine directed protect and label identify HBS-2 (K^116^, K^117^) in beta strand IX of FGF1, but do not includebeta strand VI as in FGF2[13] and HBS-3, which locates towards the N-terminus, and consists of K^24^, K^25^ and K^27^.

##### HBS-1, Arg134 and Arg137

The canonical HBS-1 is highly conserved within the FGF sub-family in terms of both sequence alignment and structure. In terms of sequence, HBS-1 in FGF1 has a high density of basic residues. The selective labelling of lysines identified Lys-127, 128, 133 and 143 as interacting with heparin. The present data demonstrate that R^134^ and R^137^ (peptides 3, 4, 5, 6, 7 and 10, Table 5; peptides 3 and 6, Table 6; Fig 6A) also bound to heparin. The involvement of R^134^ in heparin binding of FGF1was indirectly evidenced in an NMR structure of FGF1 with inositol hexasulfate as a substitute for heparin [43]. Subsequently a direct involvement was shown using a synthetic heparin hexasaccharide which caused a chemical shift perturbation of R^137^ [44,45].

**Figure 6:**
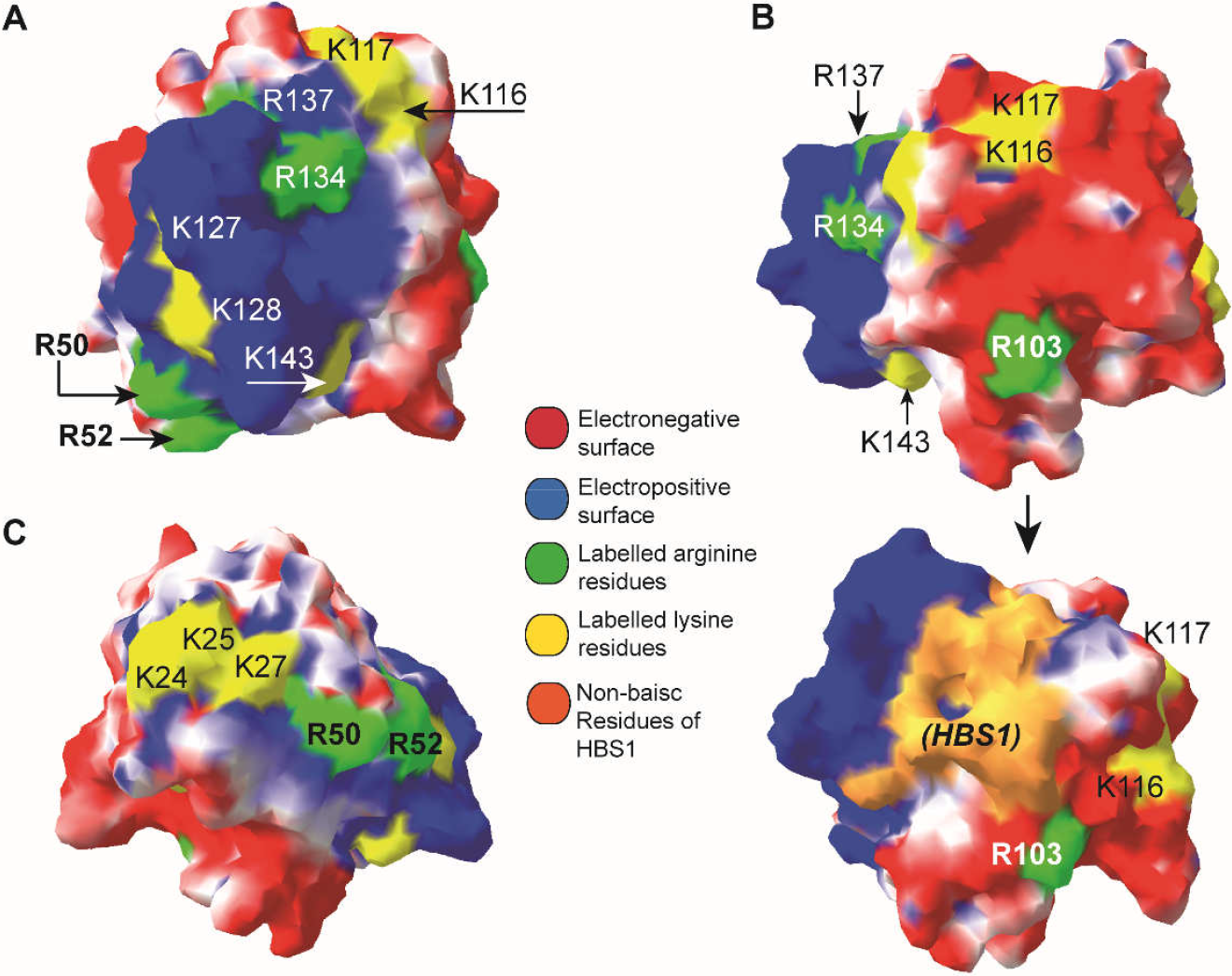
Labelled arginine and lysine residues in the heparin binding sites of FGF1. The FGF1 structure (PBD code **2erm** [44]) is shown as a surface. **A. Location of labelled R^50^, R^52^, R^134^ and R^137^ on the surface as part of the canonical HBS-1.** The other basic residues are K^127^, K^128^, K^143^, K^116^ and K^117^ [13]. K^116^ and K^117^, which previously were part of HBS-2, now are assigned to be in HBS1 by this work. **B. (Upper)** The location of is mapped to the surface, on the electronegative surface. **(Lower)** The connection of R^103^ to HBS1. **C. Locations of K^24^, K^25^ and K^27^** [13] situated at the N-terminal are shown along with locations of R^50^ and R^52^. K^24^, K^25^ and K^27^ have been reassigned from HBS-3 [13] to HBS-1 in this study.

##### FGF1 may have a single continuous HBS1

Although FGF1 has an acid pI, its charged residues are segregated on its surface so that it possesses a largely basic face and a largely acidic face (Fig 6). With the additional data provided by arginine selective labelling we were able to re-examine the assignment of the basic residues to the different HBSs.

R^103^ in beta strand VIII (Fig 7) was labelled by HPG (Tables 5, 6).On the surface of FGF1, this arginine is adjacent to HBS1 and connected to HBS-2, ^116^KK^117^ (Fig 6 B – lower panel). This suggests that R^103^ is part of HBS1. Both R^50^ and R^52^ in FGF1 were labelled by HPG, indicating that they interact with heparin. R^50^ and R^52^ locate on the loop between beta strand III and beta strand IV (Fig 7). Although separated by an acidic Asp51 residue (Fig 7), inspection of the surface electrostatic potential shows that there is no acidic boundary between these arginines (Fig 6A, C), presumably because the Asp side chain is involved in a local hydrogen bonding network. Notably, R^50^ and R^52^ are on the same basic face of the protein, as is the previously defined HBS3 (Fig 6 C). These arginines connect HBS1 and ^24^KK^25^ and K^27^ in HBS3 of FGF1 (Fig 6 C), implying that R^50^ and R^52^ as well as ^24^KK^25^ and K^27^ are part of HBS1.

**Figure 7:**
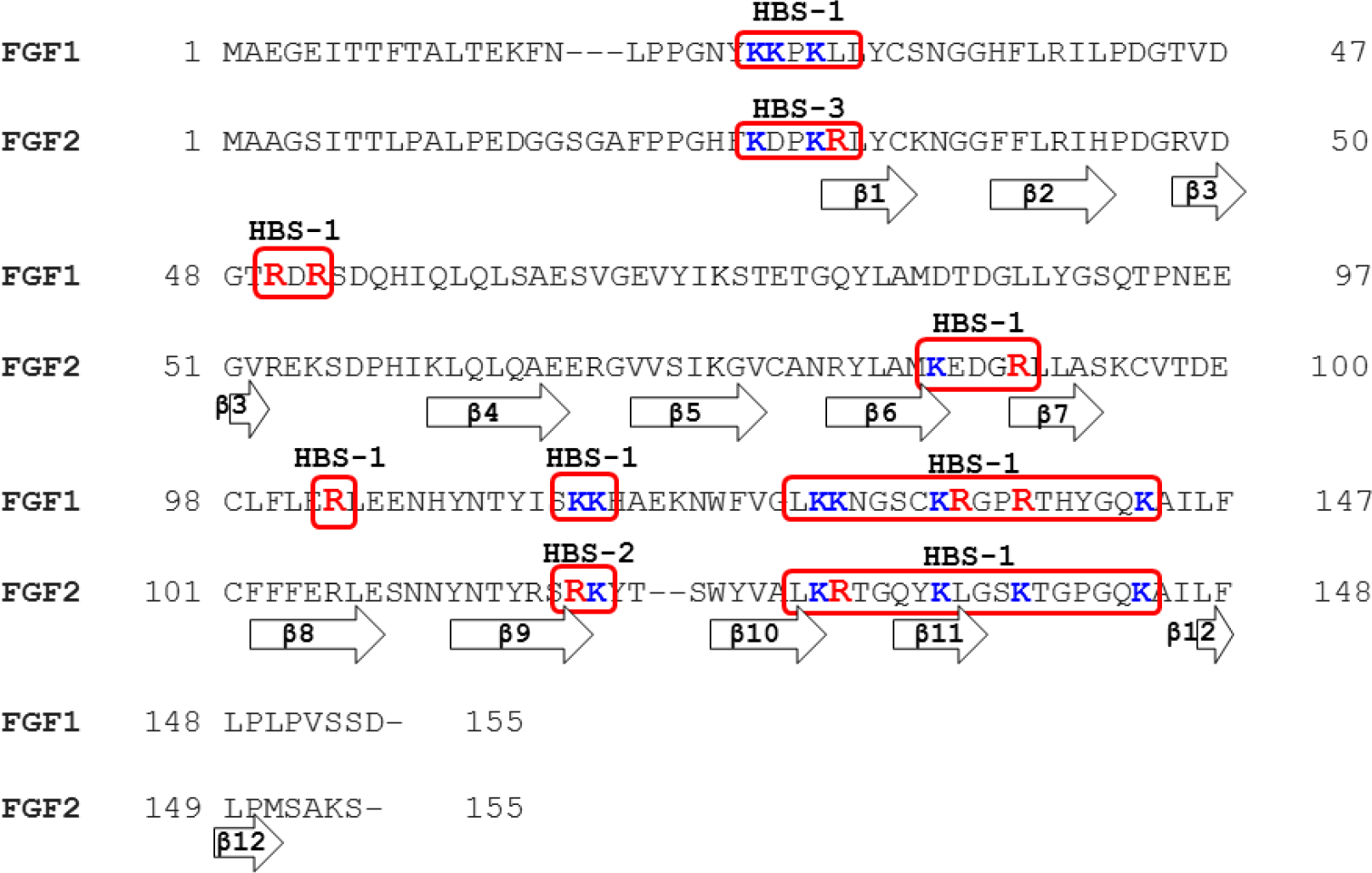
Sequence alignment of FGF2, FGF1and location of labelled arginine and lysine residues in the heparin binding sites. Uniprot ID of FGF2 sequence is, P09038-2, Uniprot ID of FGF1 sequence is P05230. Labelled arginine residues are coloured in *red*, labelled lysine residues are coloured in *blue*, the HBSs are highlighted in red boxes. The β strands are presented as the arrows.

In conclusion, we propose here that FGF1 has a single long, continuous HBS1, which comprises ^24^KK^25^ and K^27^at the N-terminus of beta strand I, R^50^ and R^52^ between β strand III/β strand IV, going through K^127^, K^128^, R^134^, R^137^, and K^143^ in β strand X/β strand XI, R^103^ in beta strand VIII and ^116^KK^117^ in beta strand IX. Thus, although the position of most of these residues in the FGF1 structure is similar to that in the structure of FGF2, differences in the surface distribution of the acidic side chains in the two proteins lead to these sites likely coalescing in FGF1.

These data demonstrate that although FGF1 and FGF2 are in the same sub-family and, possess a high level of sequence conservation, they may differ in their interactions with heparin/HS. FGF2 has the primary and the secondary binding sites separated by acidic boundaries implying that they may engage different polysaccharide chain. Indeed, this is supported by the demonstration that FGF2 can crosslink HS chains [46]. However, FGF1 possess a continuous HBS suggesting that it can bind a single HS. Analysis of the interactions of FGF1 with HS demonstrates that unlike FGF2, it does not crosslink HS chain [47]. This is consistent with FGF1 possessing a single, large HS binding site.

#### Conclusion

Our strategy for selective labelling of arginine side chains involved in binding to heparin,in combination with our published data on lysines, allows a greater dissection of the electrostatic bonds that drive the interactions between proteins and GAGs. The results with the model peptides illustrated the highly selectivity and specificity of PGO/HPG modification of the guanidino group of arginine. One of the main limitations is the multiple products from reaction with PGO, but ourstrategy of integratingmass spectrometry and automated analysis successfully tackled this issue. The data on proteins show that: 1) heparin binding effectively protects the arginine residues engagedwith the polysaccharide, allowing the protection of uninvolved arginines with PGO;2) the recovery of protein after the elution from heparin affinity mini-column is reasonably quantitative; 3) the automated analysis is sensitive enough to identify the modifications of each arginine residue. The method is rapid and so should be applicable for any protein-heparin interactions. Moreover, like lysine selective labelling, our arginine labelling method identifies the secondary, low affinity binding sites. The likely importance of these in contributing to the structure of extracellular matrix and the regulation of ligand diffusion is suggested by a recent biophysical analysis [46]. As well as protein-sulfated GAG interactions, the method should be adaptable to any interactions involving arginine residues such as protein-nucleic acid and protein-phospholipid.

The identification of arginines engaging heparin in FGF1 and FGF2 alongside the lysines involved in binding provides a number of new insights. For example, we are able to propose the reassignment of HBSs in both FGF2 and FGF1 and intriguingly, FGF1 would appear to have just a single, large HBS1, similar to FGF9 [13]. In addition, the data demonstrate that the arginine just N-terminal to β-strand I, which in at least some instances are involved in binding the cognate FGFR, can alternatively bind heparin.

## Supplementary data

**Supplementary figure 1:**
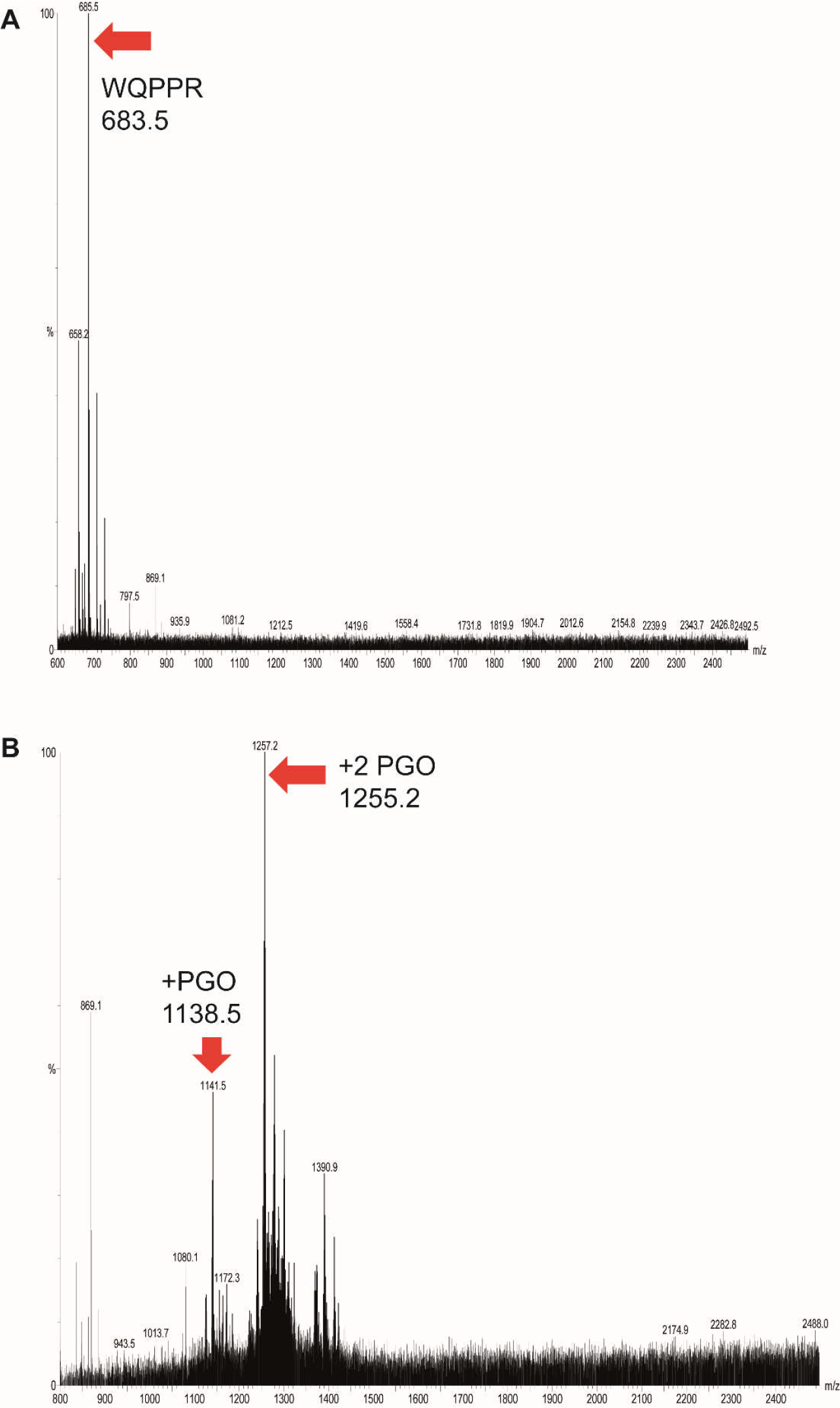
Mass spectra of the trypsin digestion products of peptide FA. **A. FA was digested by trypsin.** The larger part of the sequence was identified as “WQPPR”. B. **Peptide FA after the reaction with PGO was digested with trypsin.**

**Supplementary figure 2:**
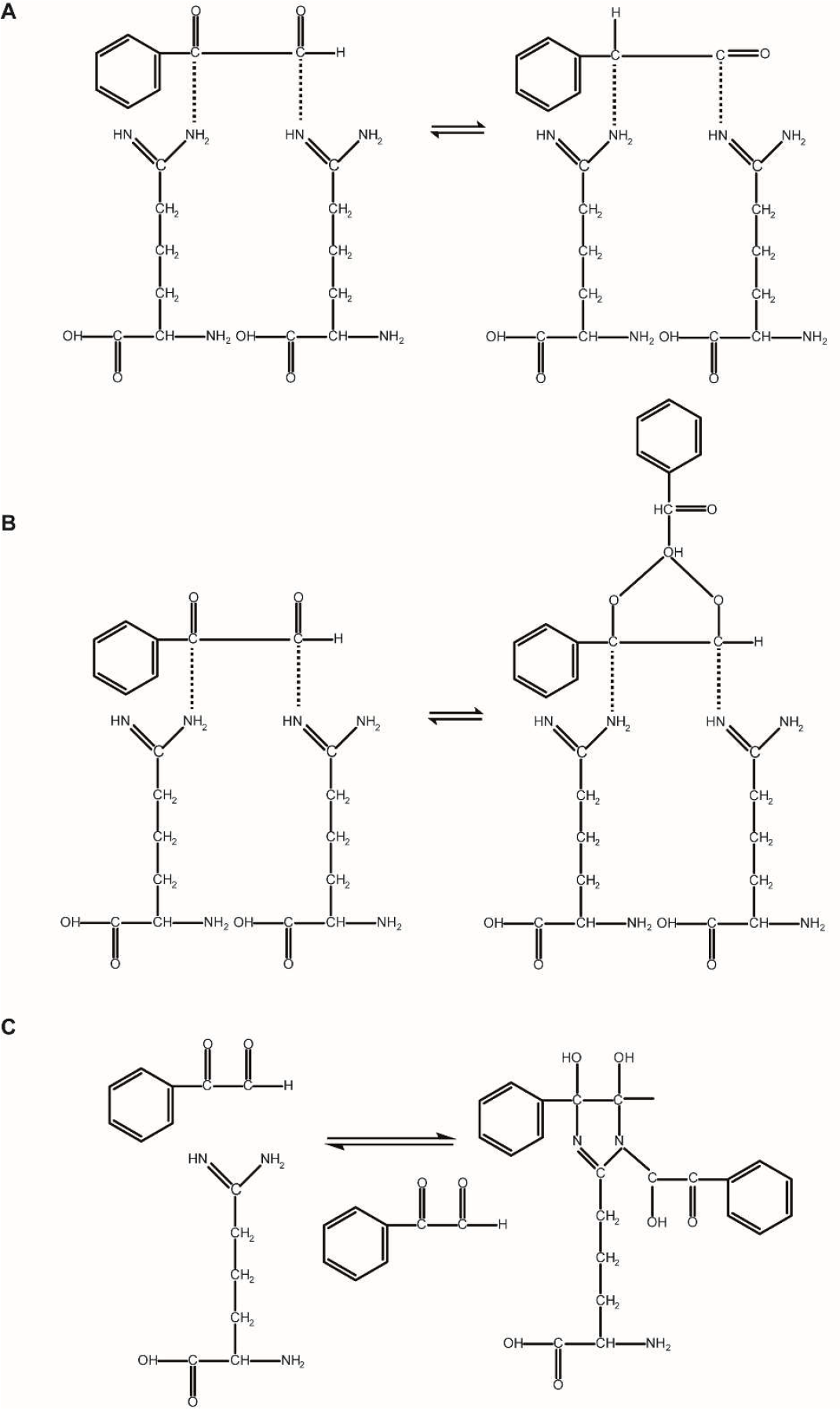
The stoichiometry of reaction of PGO with arginine residues in specific sequence contexts. A. B. PGO reacts to two adjacent arginine residues which are separated by a small sized amino acid, alanine. (A) When PGO reactsindividually with each arginine; (B) When the dicarbonyl group of PGO reacts with NH_2_ groups on different arginine residues; (C) When PGO reacts to arginine at the N- or C-terminus forming 2:1 product of PGO:Arginine.

**Supplementary figure 3:**
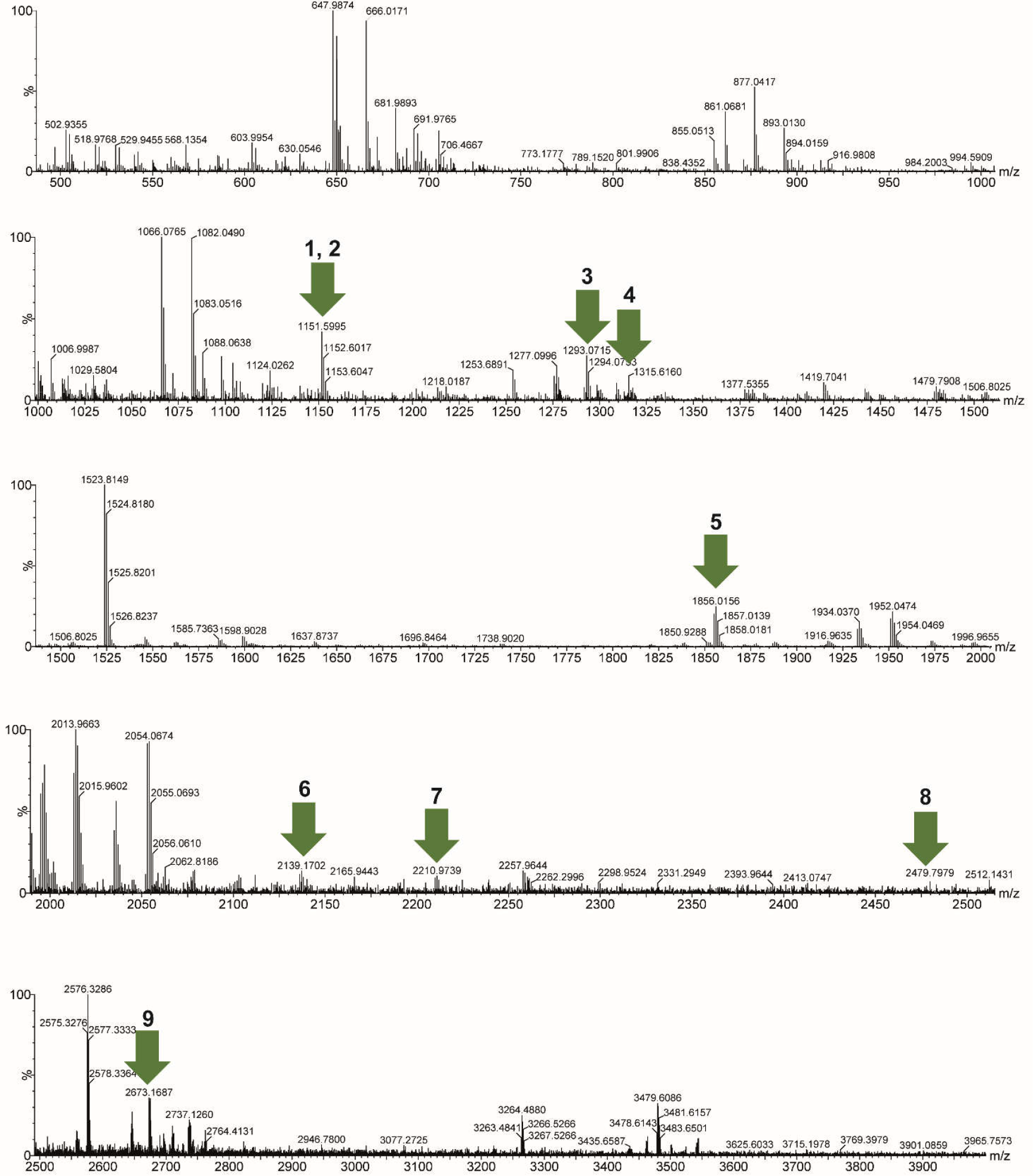
FGF2 MS spectra of peptides after in solution modification with PGO based on prediction by Prospector. The reference number of the peptide is followed by the observed m/z of the peptides produced by cleavage of FGF2 reacted in solution with PGO.

**Supplementary figure 4:**
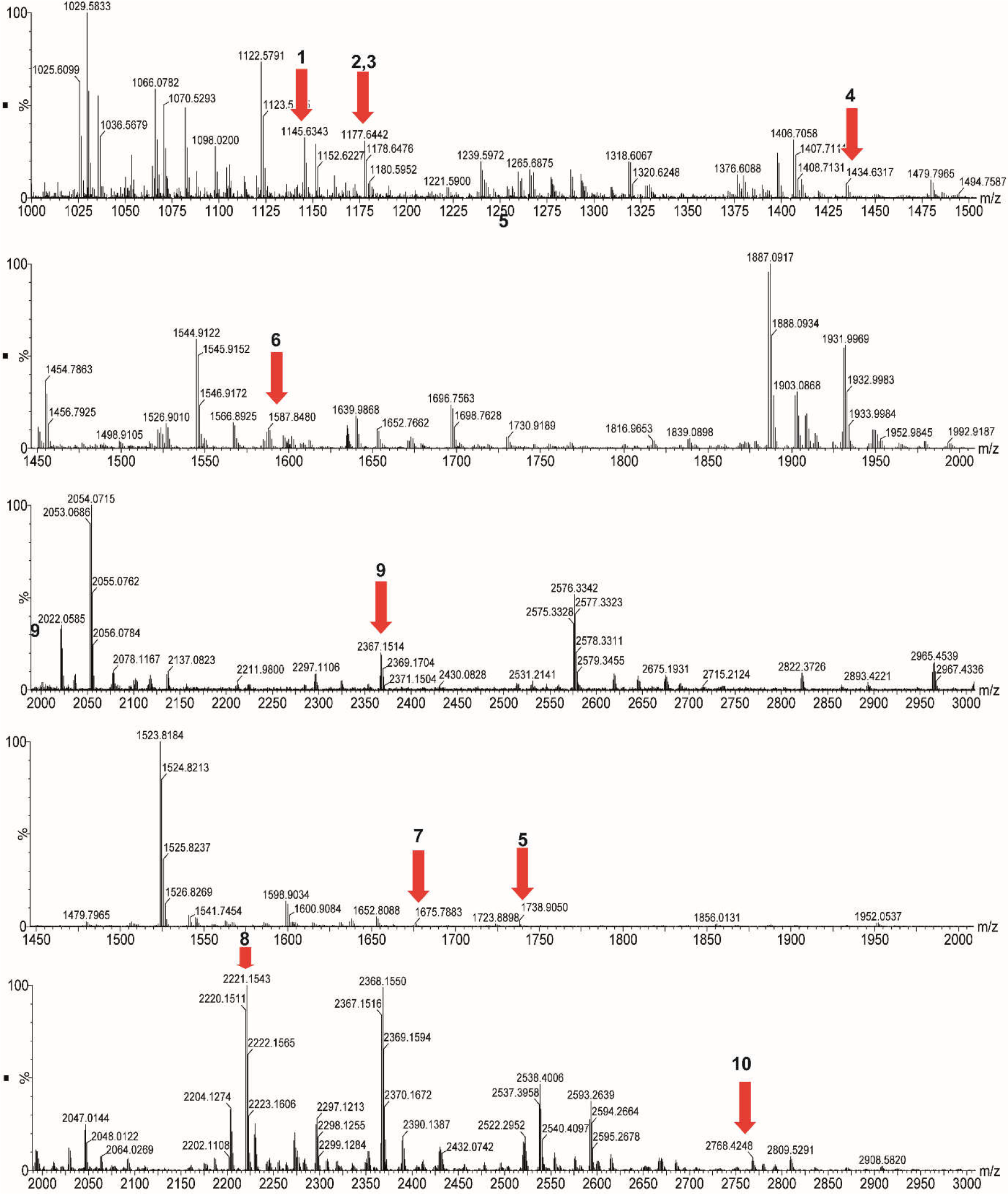
FGF2 MS spectra of peptides after modification based on prediction by Prospector. The reference number of the peptide is followed by the observed m/z of the peptides produced by the cleavage of FGF2which was before reacted with PGO when bound to a heparin affinity column (protection step) and then, following elution, reaction with HPG (labelling step).

**Supplementary figure 5:**
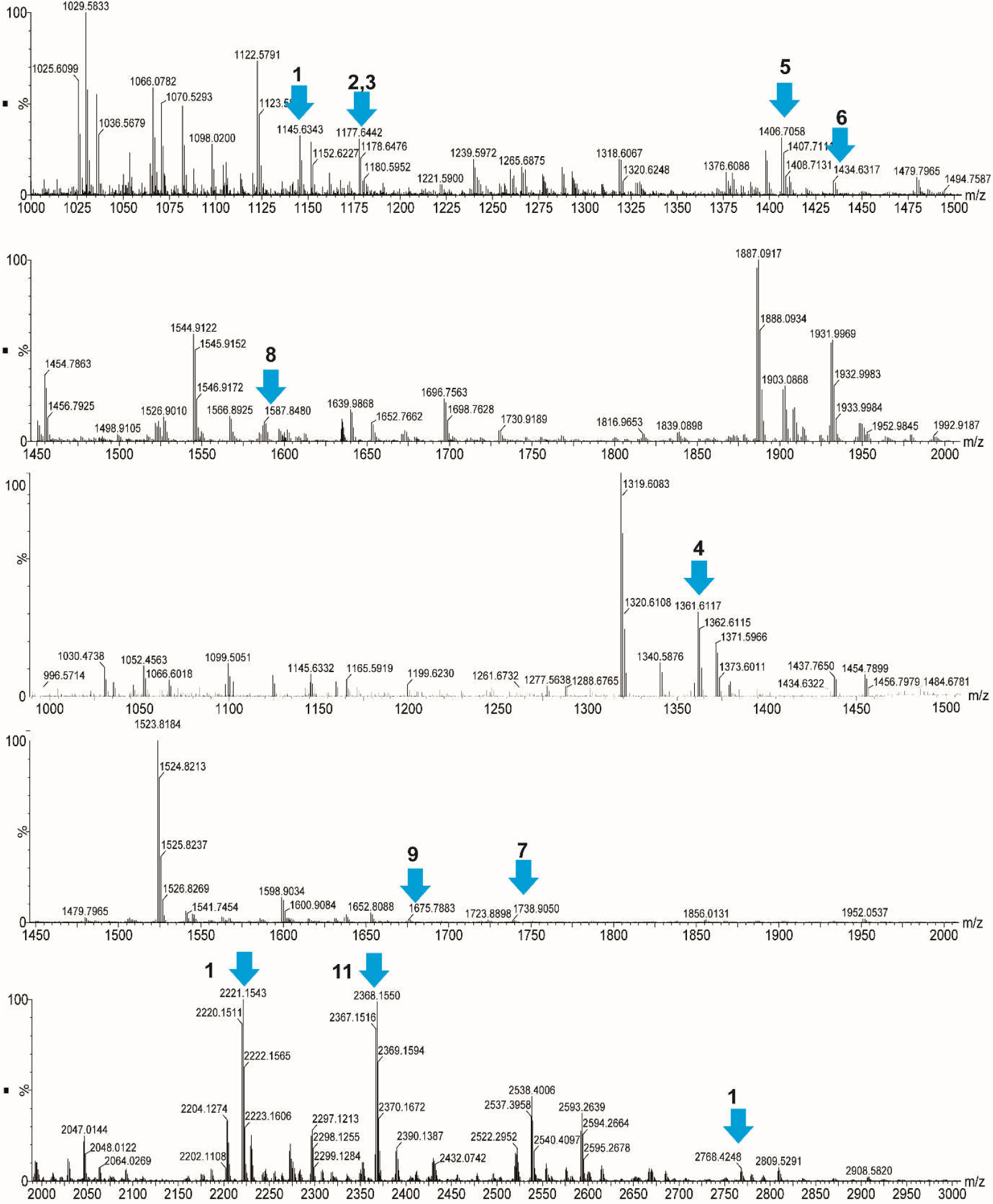
FGF2 MS spectra of peptides after modification based on prediction by Peptide Mass (ExPASy). The reference number of the peptide is followed by the observed m/z of the peptides produced by the cleavage of FGF2which was before itreacted with PGO when bound to a heparin affinity column (protection step) and then, following elution, reaction with HPG (labelling step).

**Supplementary figure 6:**
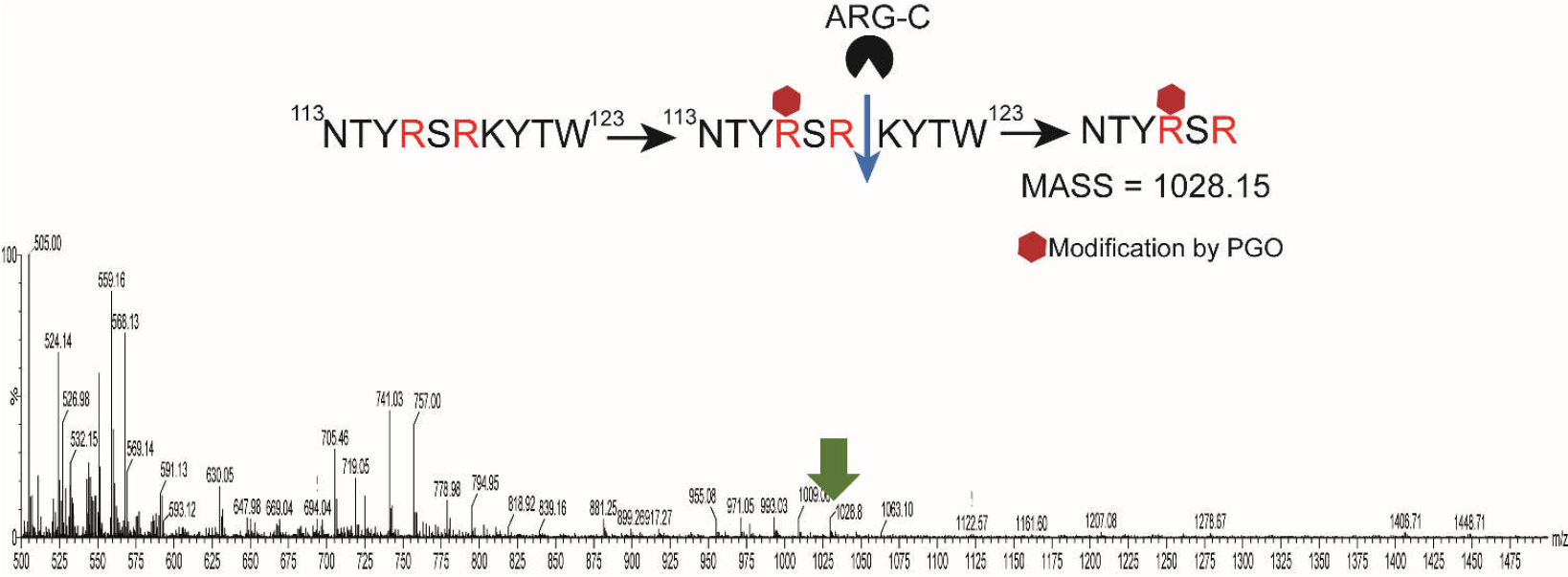
In FGF2, R^118^ engages to heparin whereas R^116^ does not. Double digestion of FGF2 by chymotrypsin and Arg-C. After the protection of exposed arginine residues on the mini-column by 200 mM PGO, protein was eluted from the columnand then cleaved for 5 hours by chymotrypsin and overnight by Arg-C. The red diamond presents for the modification by PGO on R^116^. R^118^ was cleaved by Arg-C resulting in peptide NTYRSR with the mass 1028.15, observed in the spectrum.

**Supplementary figure 7:**
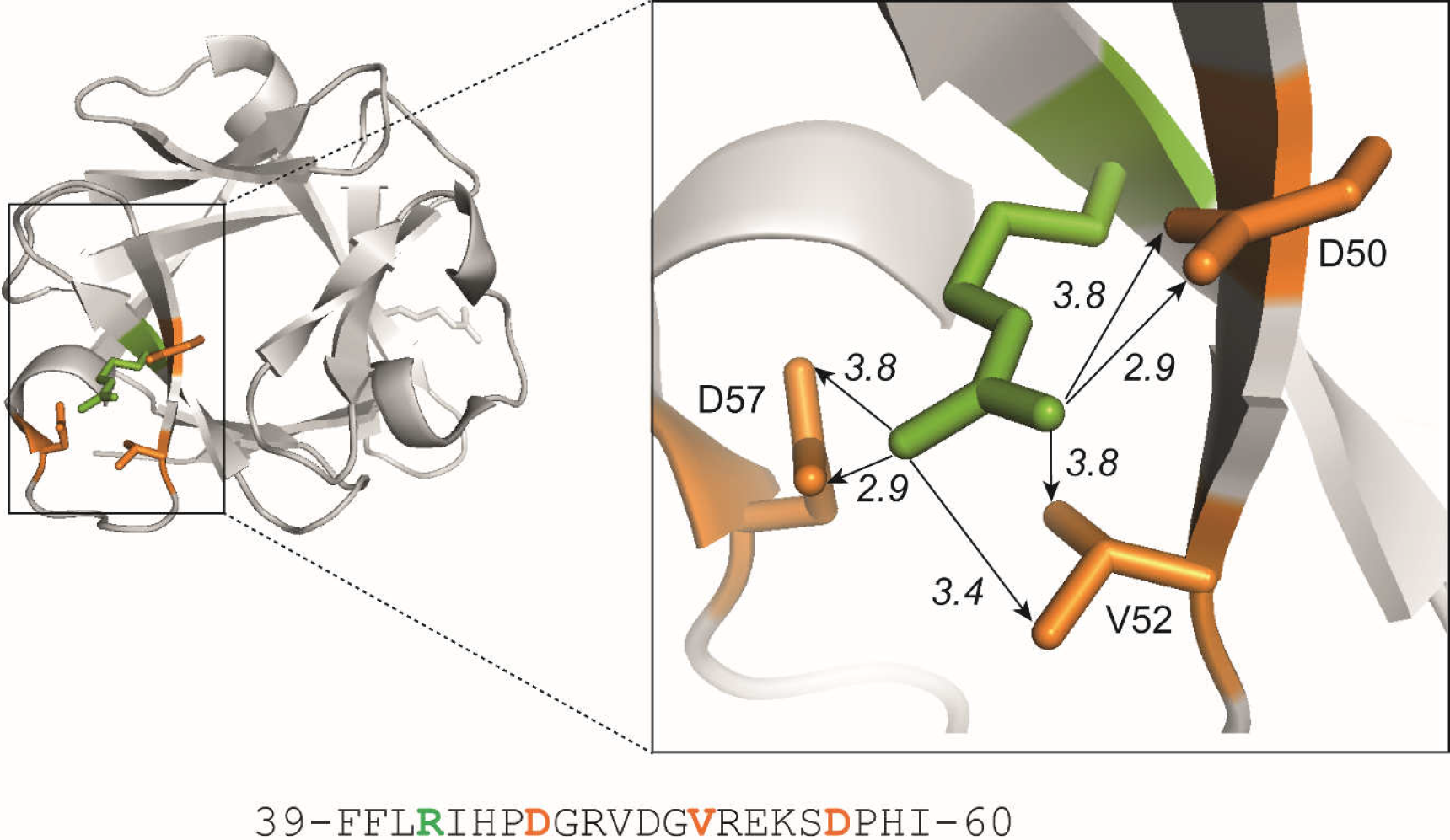
The intramolecular network of hydrogen bonds of R^42^ with D^50^, V^52^, and D^57^. The *green* presents R^42^; the *orange* shows the D^50^, V^52^, and D^57^. Their potential hydrogen bonds are presented as backlines with arrow.

**Supplementary figure 8:**
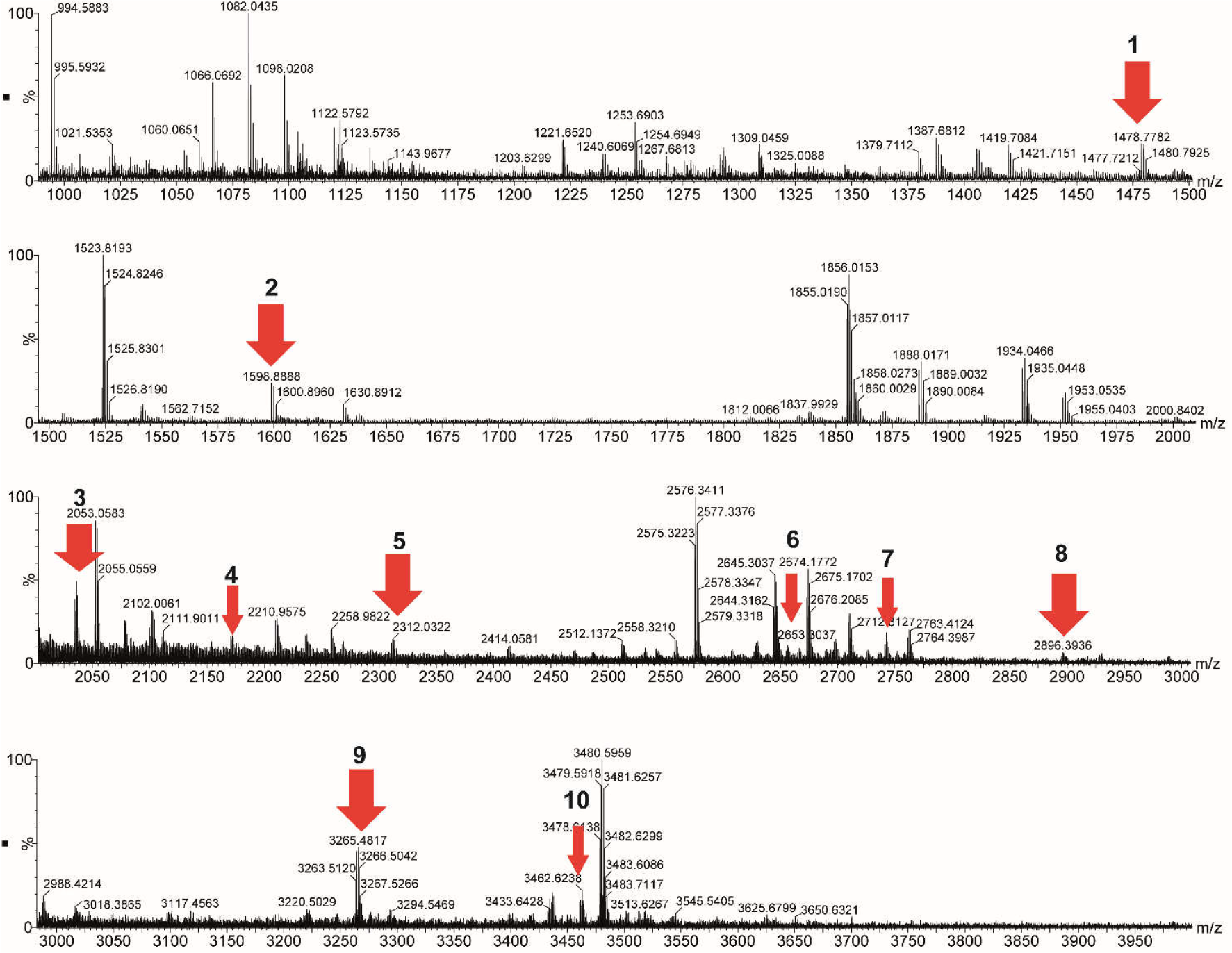
FGF1 MS spectra of peptides after modification based on prediction by Prospector. The reference number of the peptide is followed by the observed m/z of the peptides produced by the cleavage of FGF1which was before reacted with PGO when bound to a heparin affinity column (protection step) and then, following elution, reaction with HPG (labelling step).

**Supplementary figure 9:**
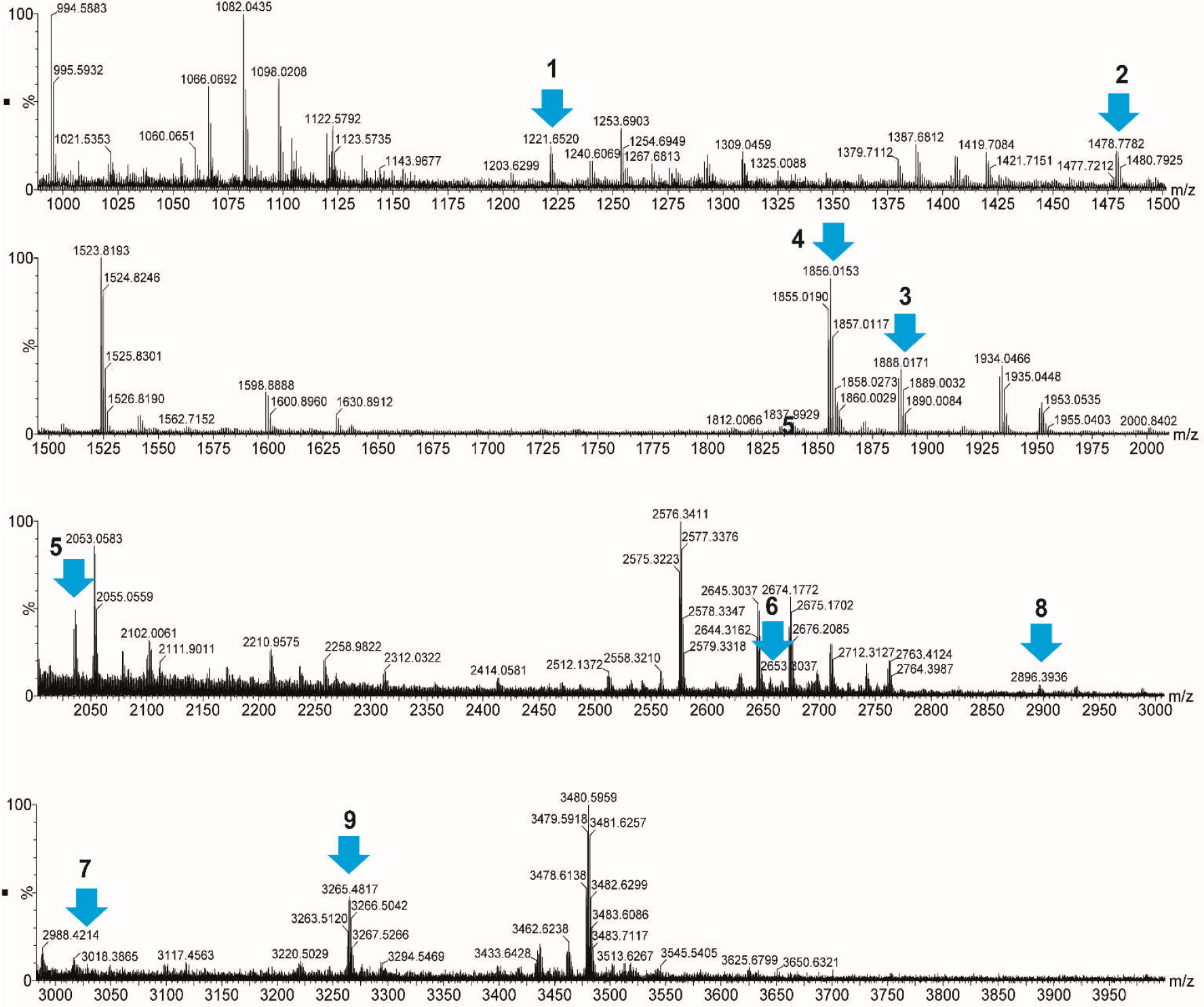
FGF1 MS spectra of peptides after modification based on prediction by Peptide Mass (ExPASy). The reference number of the peptide is followed by the observed m/z of the peptides produced by the cleavage of FGF1which was before reacted with PGO when bound to a heparin affinity column (protection step) and then, following elution, reaction with HPG (labelling step).

**Supplementary table 1:**
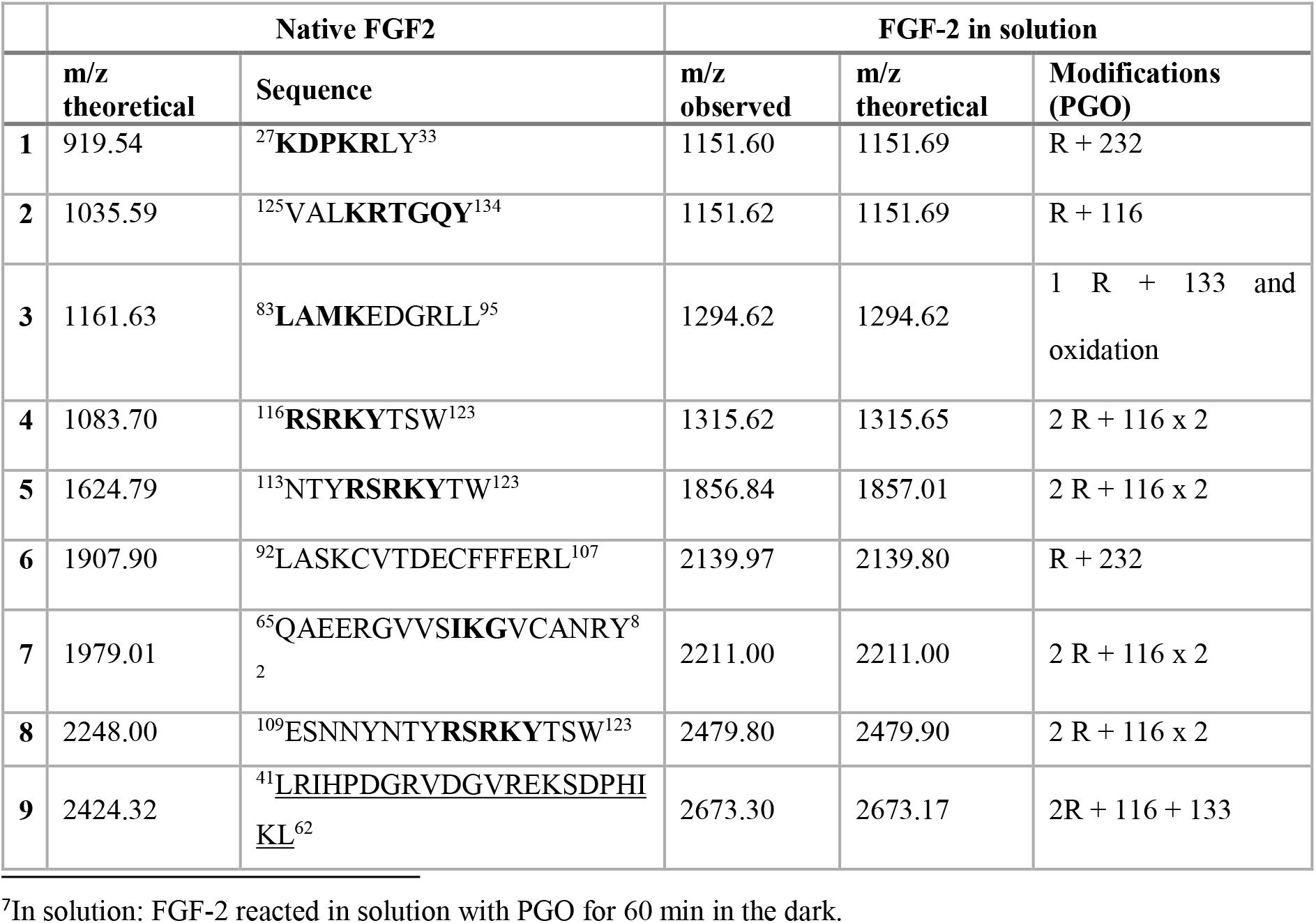
FGF-2 MS analysis based on prediction by Prospector. The reference number of the peptide is followed by the predicted m/z and with the sequence of the peptides following cleavage of native FGF-2 chymotrypsin. The two columns under “Native FGF-2” present the predicted m/z and sequences of peptides after chymotrypsin digestion of FGF-2.The first two columns under “FGF-2 in-solution” present the observed and predicted m/z for FGF-2 after modification by PGO^7^. The final column indicates the modification occurring on the arginine residues of the peptides.

**Supplementary table 2:**
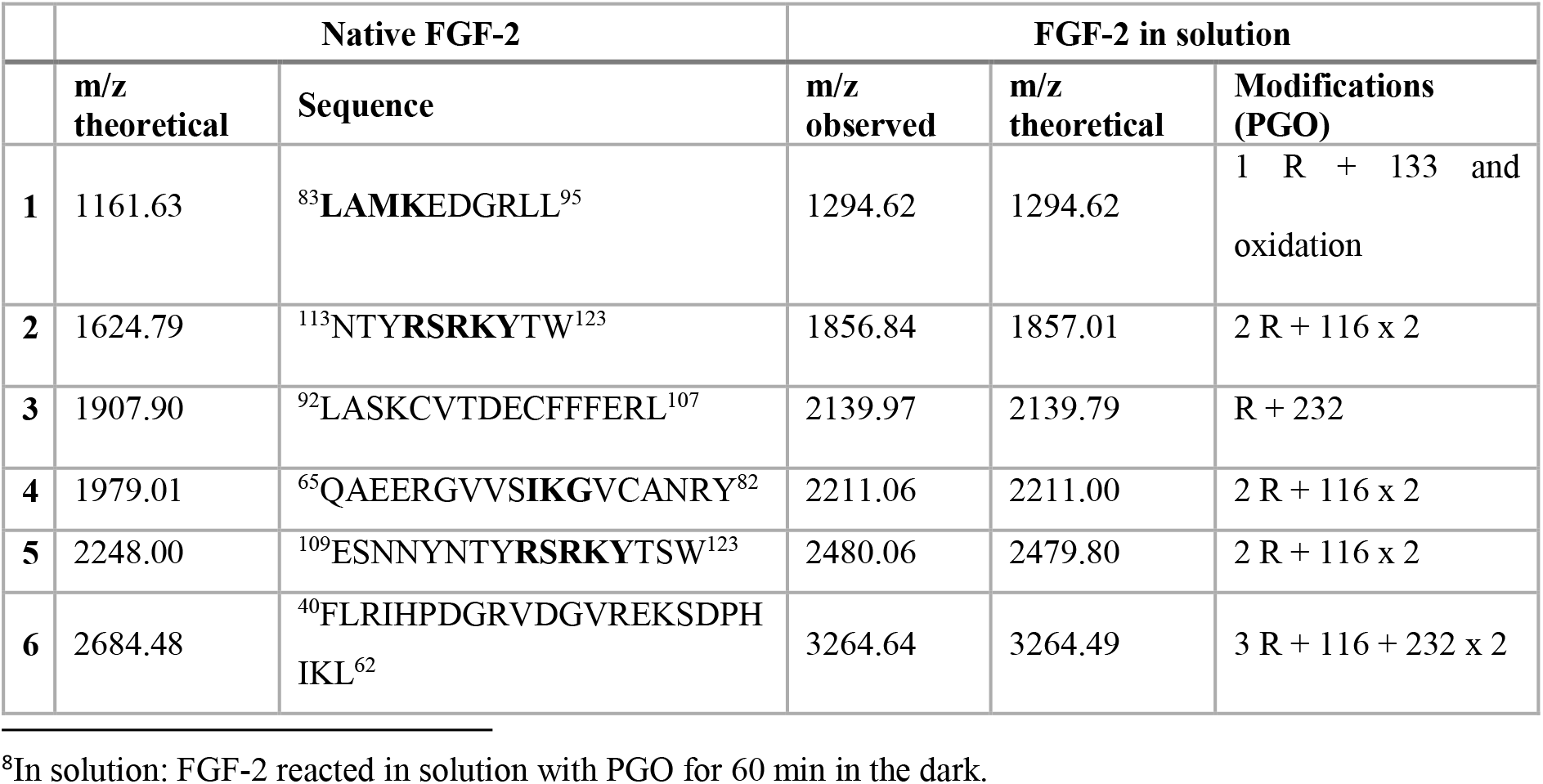
FGF-2 MS analysis based on prediction by Peptide Mass (ExPASY) The reference number of the peptide is followed by the predicted m/z and with the sequence of the peptides following cleavage of native FGF-2 chymotrypsin. The two columns under “Native FGF-2” present the predicted m/z and sequences of peptides after chymotrypsin digestion of FGF-2.The first two columns under “FGF-2 in-solution” present the observed and predicted m/z for FGF-2 after modification by PGO^8^. The final column indicates the modification occurring on the arginine residues of the peptides.

## References

1. Dreyfuss JL, Regatieri CV., Jarrouge TR, Cavalheiro RP, Sampaio LO, Nader HB. Heparan sulfate proteoglycans: structure, protein interactions and cell signaling. An Acad Bras Cienc. 2009;81: 409–429. doi:10.1590/S0001-37652009000300007

2. Ori A, Wilkinson MC, Fernig DG. The heparanome and regulation of cell function: structures, functions and challenges. Front Biosci. 2008; 4309. doi:10.2741/3007

3. Duchesne L, Octeau V, Bearon RN, Beckett A, Prior IA, Lounis B, et al. Transport of fibroblast growth factor 2 in the pericellular matrix is controlled by the spatial distribution of its binding sites in Heparan Sulfate. PLoS Biol. 2012;10: e1001361. doi:10.1371/journal.pbio.1001361

4. Fernig DG, Mason D, Marcello M, Li Y, Sun C, Lévy R. Selectivity in glycosaminoglycan binding dictates the distribution and diffusion of fibroblast growth factors in the pericellular matrix. Open Biol. 2016;6: 150277. doi:10.1098/rsob.150277

5. Olsen SK, Ibrahimi OA, Raucci A, Zhang F, Eliseenkova A V, Yayon A, et al. Insights into the molecular basis for fibroblast growth factor receptor autoinhibition and ligand-binding promiscuity. PNAS. 2004;101: 935–940.

6. Schlessinger J, Plotnikov AN, Ibrahimi OA, Eliseenkova A V., Yeh BK, Yayon A, et al. Crystal structure of a ternary FGF-FGFR-heparin complex reveals a dual role for heparin in FGFR binding and dimerization. Mol Cell. 2000;6: 743–750. doi:10.1016/S1097-2765(00)00073-3

7. Dementiev A, Petitou M, Herbert J-M, Gettins PGW. The ternary complex of antithrombin–anhydrothrombin–heparin reveals the basis of inhibitor specificity. Nat Struct Mol Biol. 2004;11: 863–867. doi:10.1038/nsmb810

8. Fromm JR, Hileman RE, Caldwell EE, Weiler JM, Linhardt RJ. Differences in the interaction of heparin with arginine and lysine and the importance of these basic amino acids in the binding of heparin to acidic fibroblast growth factor. Arch Biochem Biophys. 1995;323: 279–87. doi:10.1006/abbi.1995.9963

9. Olson ST, Halvorson HR, Bjork I. Quantitative characterization of the thrombin-heparin interaction: Discrimination between specific and nonspecific binding models. J Biol Chem. 1991;266: 6342–6352.

10. Rousselle P, Vivès RR, Crublet E, Lortat-Jacob H, Gagnon J, Andrieu J-P. A novel strategy for defining critical amino acid residues involved in protein/glycosaminoglycan interactions. J Biol Chem. 2004;279: 54327–54333. doi:10.1074/jbc.m409760200

11. Ori A, Free P, Courty J, Wilkinson MC, Fernig DG. Identification of heparin-binding sites in proteins by selective labeling. Mol Cell Proteomics. 2009;8: 2256–2265. doi:10.1074/mcp.M900031-MCP200

12. Li Y, Sun C, Yates EA, Jiang C, Wilkinson MC, Fernig DG. Heparin binding preference and structures in the fibroblast growth factor family parallel their evolutionary diversification. Open Biol. 2016;6.

13. Xu R, Ori A, Rudd TR, Uniewicz KA, Ahmed YA, Guimond SE, et al. Diversification of the structural determinants of fibroblast growth factor-heparin interactions: Implications for binding specificity. J Biol Chem. 2012;287: 40061–40073. doi:10.1074/jbc.M112.398826

14. Migliorini E, Thakar D, Ku J, Sadir R, Dyer DP, Li Y, et al. Cytokines and growth factors cross-link heparan sulfate. Open Biol. 2015;

15. Mayo KH, Ilyina E, Roongta V, Dundas M, Joseph J, Lai CK, et al. Heparin binding to platelet factor-4. An NMR and site-directed mutagenesis study: arginine residues are crucial for binding. Biochem J. 1995;312

16. Moy FJ, Safran M, Seddon AP, Kitchen D, Böhlen P, Aviezer D, et al. Properly oriented heparin-decasaccharide-induced dimers are the biologically active form of basic fibroblast growth factor. Biochemistry. 1997;36: 4782–4791. doi:10.1021/bi9625455

17. Thompson LD, Pantoliano MW, Springer BA. Energetic characterization of the basic fibroblast growth factor-heparin interaction: identification of the heparin binding domain. Biochemistry. 1994;33: 3831–3840. doi:10.1021/bi00179a006

18. Springer BA, Pantoliano MW, Barbera FA, Gunyuzlu PL, Thompson LD, Herblin WF, et al. Identification and concerted function of 2 receptor-binding surfaces on basic fibroblast growth-factor required for mitogenesis. J Biol Chem. 1994;269: 26879–26884.

19. Amara A, Lorthioir O, Magerus A, Thelen M, Montes M, Virelizier J, et al. Stromal cell-derived factor-1 α associates with heparan sulfates through the first β-strand of the chemokine. J Biol Chem. 1999;274: 23916–23925. doi:10.1074/jbc.274.34.23916

20. Faham S, Hileman RE, Fromm JR, Linhardt RJ, Rees DC. Heparin Structure and Interactions with Basic Fibroblast Growth Factor. Science (80). 1996;271: 1116–1120. doi:10.1126/science.271.5252.1116

21. Lee J, Wee S, Gunaratne J, Chua RJE, Smith RAA, Ling L, et al. Structural determinants of heparin-transforming growth factor-β1 interactions and their effects on signaling. Glycobiology. 2015;25: 1491–1504. doi:10.1093/glycob/cwv064

22. Dos Santos C, Blanc C, Elahouel R, Prescott M, Carpentier G, Ori A, et al. Proliferation and migration activities of fibroblast growth factor-2 in endothelial cells are modulated by its direct interaction with heparin affin regulatory peptide. Biochimie. 2014;107: 350–357. doi:10.1016/j.biochi.2014.10.002

23. Mendoza VL, Vachet RW. Probing Protein Structure by Amino Acid-Specific Covalent Labeling and Mass Spectrometry Vanessa. Mass Spectrom Rev. 2010;28: 785–815. doi:10.1002/mas.20203.

24. Takahashi K. Chemistry and metabolism of macromolecules: The reaction of phenylglyoxal with arginine residues in proteins. J Biol Chem. 1968;243: 6171–6179. Available: http://www.jbc.org/content/243/23/6171.

25. Morkin E, Flink IL, Surath K Banerjee. Phenylglyoxal modification of Cardiac myosin S-l: Evidence for essential arginine residues at the active sites. J Biol Chem. 1979;254: 12647–12652.

26. Vanoni MA, Simonetta MP, Curti B, Negri A, Ronchi S. Phenylglyoxal modification of arginines in mammalian D-amino-acid oxidase. Eur J Biochem. 1987;167: 261–7A.

27. Poulose AJ, P. E. Kolattukudy. Enzymatic reduction of phenylglyoxal and 2, 3-butanedione, two commonly used arginine-modifying reagents, by the ketoacyl reductase domain of fatty acid synthase. Int J Biochem. 2001;18: 807–812.

28. Pandeswari PB, Sabareesh V, Vijayalakshmi MA. Insights into stoichiometry of arginine modification by phenylglyoxal and 1, 2-cyclohexanedione probed by LC-ESI-MS. J Proteins Proteomics. 2016;7: 323–347.

29. Uniewicz K, Ori A, Xu R, Ahmed Y, Fernig DG, Yates E. Differential Scanning Fluorimetry measurement of protein stability changes upon binding to glycosaminoglycans: a rapid screening test for binding specificity. Anal Chem. 2010;82: 1–3.

30. Chevallet M, Luche S, Rabilloud T. Silver staining of proteins in polyacrylamide gels. Nat Protoc. 2006;1: 1852–1858. doi:10.1038/nprot.2006.288

31. Takahashp K. Further Studies on the Reactions of Phenylglyoxal and related reagents with proteins. J Biol Chem. 1977;414: 403–414.

32. Shu-Tong Cheung and Margaret L. Fonda. Reaction of phenylglyoxal with arginine: the effect of buffers and pH. Biochem Biophys Res Commun. 1979;90: 940–947.

33. Eriksson O, Fontaine E, Bernardi P. Chemical modification of arginines by 2,3-butanedione and phenylglyoxal causes closure of the mitochondrial permeability transition pore. J Biol Chem. 1998;273: 12669–12674. doi:10.1074/jbc.273.20.12669

34. Wood TD, Guan Z, Borders CL, Chen LH, Kenyon GL, McLafferty FW. Creatine kinase: essential arginine residues at the nucleotide binding site identified by chemical modification and high-resolution tandem mass spectrometry. Proc Natl Acad Sci U S A. 1998;95: 3362–5. doi:10.1073/pnas.95.7.3362

35. Junkova P, Vermachova M, Prchal J, Kuckova S, Hrabal R, Hynek R, et al. Improved approach for the labeling of arginine, glutamic, and aspartic acid side chains in proteins using chromatographic techniques. J Liq Chromatogr Relat Technol. 2013;36: 1221–1230. doi:10.1080/10826076.2012.685918

36. Ibrahimi OA, Yeh BK, Eliseenkova A V., Zhang F, Olsen SK, Igarashi M, et al. Analysis of mutations in Fibroblast Growth Factor (FGF) and a pathogenic mutation in FGF Receptor (FGFR) provides direct evidence for the symmetric two-end model for FGFR dimerization. Mol Cell Biol. 2005;25: 671–684. doi:10.1128/MCB.25.2.671-684.2005

37. Pellegrini L, Burke DF, von Delft F, Mulloy B, Blundell TL. Crystal structure of fibroblast growth factor receptor ectodomain bound to ligand and heparin. Nature. 2000;407: 1029–34. doi:10.1038/35039551

38. Baird A, Schubert D, Ling N, Guillemin R. Receptor- and heparin-binding domains of basic fibroblast growth factor. Proc Natl Acad Sci U S A. 1988;85: 2324–2328. doi:10.1073/pnas.85.7.2324

39. Plotnikov AN, Schlessinger J, Hubbard SR, Mohammadi M. Structural basis for FGF receptor dimerization and activation. Cell. 1999;98: 641–650. doi:10.1016/S0092-8674(00)80051-3

40. Aviezer D, Safran M, Yayon A. Heparin differentially regulates the interaction of fibroblast growth factor-4 with FGF receptors 1 and 2. Biochem Biophys Res Commun. 1999;263: 621–626. doi:10.1006/bbrc.1999.1434

41. Lozano RM, Pineda-Lucena A, Gonzalez C, Jiménez MÁ, Cuevas P, Redondo-Horcajo M, et al. ^1^H NMR structural characterization of a nonmitogenic, vasodilatory, ischemia-protector and neuromodulatory acidic fibroblast growth factor. Biochemistry. 2000;39: 4982–4993. doi:10.1021/bi992544n

42. Mach H, Volkin DB, Burke CJ, Middaugh CR, Mattsson L. Nature of the interaction of heparn with acidic fibroblast growth factor. Biochemistry. 1993;32: 5480–5489.

43. A. Pineda-Lucena, M. A. Jimezez, J. L. Nieto, J. Santoro MR and GG-G. H-NMR assignment and solution structure of Human acidic Fibroblast growth factor activated by Inositol Hexasulfate. 1994. pp. 81–98.

44. Canales A, Lozano R, López-Méndez B, Angulo J, Ojeda R, Nieto PM, et al. Solution NMR structure of a human FGF-1 monomer, activated by a hexasaccharide heparin-analogue. FEBS J. 2006;273: 4716–4727. doi:10.1111/j.1742-4658.2006.05474.x

45. Ogura K, Nagata K, Hatanaka H, Habuchi H, Kimata K, Tate S, et al. Solution Structure of Human Acidic Fibroblast Growth Factor and Interaction with Heparin-Derived Hexasaccharide. J Biomol NMR. 1999;13: 11–24. doi:10.1023/A:1008330622467

46. Migliorini E, Thakar D, Kühnle J, Sadir R, Dyer DP, Li Y, et al. Cytokines and growth factors cross-link heparan sulfate. Open Biol. 2015;5. doi:10.1098/rsob.15.0046

47. Baradji Aïseta. Interactions of fibroblast growth factors with glycosaminoglycan brushes and the pericellular matrix. Thesis. 2017. doi:10.1016/j.injury.2012.08.014

48. Ago H, Kitagawa Y, Fujishima A, Matsuura Y, Katsube Y. Crystal structure of basic fibroblast growth factor at 1.6 Angstron resolution. J Biochem. 1991;110: 360–363. doi:10.1093/oxfordjournals.jbchem.a123586

